# Nanoplasmonic SERS-enabled structural identification of an uncharacterised indole metabolite directly in E. coli cultures

**DOI:** 10.1101/2025.11.10.687708

**Authors:** Mo Vali, Elle Wyatt, Kieran Abbott, Cameron Croft, Pietro Lio, Thomas F. Krauss, Minahil Khan, Ashraf Zarkan, Jeremy J. Baumberg, Diana Fusco

**Affiliations:** Cavendish Laboratory, Department of Physics, University of Cambridge, JJ Thomson Avenue, Cambridge, UK; Department of Genetics, University of Cambridge, Cambridge, UK; Department of Computer Science and Technology, University of Cambridge, Cambridge, UK; Department of Physics, Engineering and Technology, University of York, York, UK

**Keywords:** SERS, nanoplasmonic sensing, metabolite, genetics, bacteria, indole, structural identification, signalling molecules

## Abstract

Tryptophanase (TnaA) is a promiscuous enzyme which regulates the production of amino acid-derived metabolites, essential for bacterial communication. TnaA is known to convert tryptophan into indole, an important signalling molecule influencing a range of behaviours - from bacterial quorum sensing to urinary tract infections (UTI) - but other conversion routes remain obscure. Here, we show that nanoplasmonic surface-enhanced Raman spectroscopy (SERS) combined with targeted genetic subtraction enables the label-free, cellular characterisation and structural identification of TnaA-derived metabolites directly from *Escherichia coli* culture solutions, without isolation or purification. We systematically profile the SERS spectra of wild-type and *tnaA* gene knockout strains (*E. coli* BW25113 and the uropathogenic 536) supplemented with each of the twenty amino acids, uncovering a previously unreported TnaA-dependent metabolic signature, I*, whose Raman fingerprint does not match free indole or any common indole derivative characterised by mass spectrometry. Using isotopomer-labelled substrates together with structural reference compounds, we perform label-free structural deduction of I* directly from SERS spectra. We find I*’s structure is consistent with an intact indole ring carrying a C3 substituent, likely produced indirectly from intracellular tryptophan. These findings introduce a novel indole metabolite, raise new hypotheses regarding the biological role of TnaA-mediated byproducts in bacterial signalling and virulence, and demonstrate that nanoplasmonic SERS provides a powerful framework for probing enzyme activity and bioactive metabolite production directly from living bacterial cultures with nanomolar sensitivity and structural specificity.

## Introduction

Bacteria produce a wide chemical portfolio of bioactive extracellular molecules that regulate behavioural phenotypes, cell-to-cell communication and response to environmental cues, as well as biofilm formation [1–6]. Many of these metabolites have also been linked to infections, inflammation, neurological disorders and changes in the human microbiome [7–11]. These molecules have attracted significant research interest, as their production can be engineered for a wide range of uses, from antibiotics to fertiliser agents [12, 13]. *In vivo* measurements of metabolites are thus essential both to better understand their role in bacterial physiology in the wild and to harness their production for practical applications.

Indole is an important example of such signalling molecules produced by over 85 Gram-positive and Gram-negative bacterial species[14–17] and known to cause a wide range of effects on membrane potential, cytoplasmic pH, transport, biofilm formation, and antibiotic resistance and persistence [16]. Indole is clinically relevant, acting as a biochemical precursor to neurotransmitters such as serotonin, and frequently detected in infections caused by *E. coli*, particularly in the urinary tract [18–21]. Indole is produced by the tetrameric enzyme tryptophanase, or tryptophan indole-lyase, EC 4.1.99.1 (TnaA), which catalyses *α, β*-elimination and *β*-substitution reactions of both natural and synthetic amino acids. The *tnaA* gene encoding the enzyme TnaA has been reported in over 190 species, highlighting the pervasive role of TnaA in metabolism[22].

It is well established that TnaA catalyses the conversion of L-tryptophan to indole, pyruvate (via the removal of the carboxyl group), and ammonia (via removal of the amino group) [23]. However, the role of TnaA in other metabolic pathways remains poorly characterised. Seminal work by Newton and Snell in 1964 reported that crystalline preparations of TnaA could catalyse a series of *α, β*-elimination and *β*-substitution reactions in amino acids beyond tryptophan [24, 25]. Subsequent work has utilised spectrophotometry techniques to identify coenzyme-substrate intermediates using characteristic absorption and circular dichroic spectra [26–29], confirming that other amino acids beyond tryptophan can act as substrates (and inhibitors) of TnaA *in vitro* [26]. Few studies have systematically evaluated the role of TnaA, and to our knowledge, there is no study that has carried out a comprehensive comparison of the *in vitro* enzymatic activity of TnaA versus its *in vivo* cellular counterpart. Such studies are needed to fully understand the TnaA-regulated metabolic pathways in the living cell, and their role in indole production.

Conventional microbial typing and diagnostic approaches for detecting metabolic byproducts or bioactive metabolites typically require extensive sample preparation, often relying on specific reagents or fluorescent markers. Surface-enhanced Raman Spectroscopy (SERS) is becoming increasingly employed as an alternative technology in microbiological contexts, as it has shown high sensitivity for label- and stain-free bacterial detection, identification, and antimicrobial susceptibility testing, both on solid and liquid samples [30–33]. In contrast to conventional microbial typing methods, SERS has demonstrated superior sensitivity (10^−9^M to 10^−12^M) and is largely agnostic to the target metabolite, as long as the Raman cross-section is sufficiently large [34]. SERS capabilities have been illustrated for the identification of taxonomically-close bacterial species [35, 36], chemical profiling of microbial cells [37, 38], single cell analysis [39] and *in vivo* diagnostics [35, 40, 41]. So far, only a limited number of studies have employed SERS for characterising bacterial metabolite profiles, as label-free detection of target analytes in complex biofluids still presents considerable technical challenges in isolating interpretable chemical signatures. Notably, Bodelon et al. developed a method for label-free SERS detection of pyocyanin, a proxy analyte for quorum sensing, in *Pseudomonas aeruginosa* and *Chromobacterium violaceum* biofilms. The authors made use of specially designed nanostructured hybrid materials, demonstrating the potential of SERS to track a known target molecule in bacterial aggregates [42]. Nevertheless, to our knowledge, SERS has not been utilised to characterise an enzyme’s metabolic activity directly *in vivo* or to identify its primary byproducts without specific extraction.

In recent work, we have demonstrated the construction and use of precision nanogaps within sheets of self-assembled gold nanoparticles as reproducible and re-useable SERS devices for sensing bioanalytes such as neurotransmitters [43–45]. These ‘MLagg’ films consist of a near-monolayer aggregate of close-packed gold nanoparticles (here 80nm diameter), with 0.9 nm particle spacing precisely defined by the rigid cucurbit[n]uril (CB[n]) spacer molecule employed (Fig. 1a). Pre-cleaning these nanogaps to remove all organics inherent in their self-assembly yields precision SERS devices that can be recleaned of all analytes and fouling for reuse. The nanogaps give very large plasmonic SERS enhancements (*>* 10^8^) for small molecules adsorbing within the plasmonic hotspots in the gaps between the nanoparticles, enabling the detection of very low (nM) molecular concentrations. The consistent preparation of these MLaggs and their reproducible response opens up the capability to study many biomolecules in complex environments over long periods of time, affordably and sustainably.

**Figure 1.**
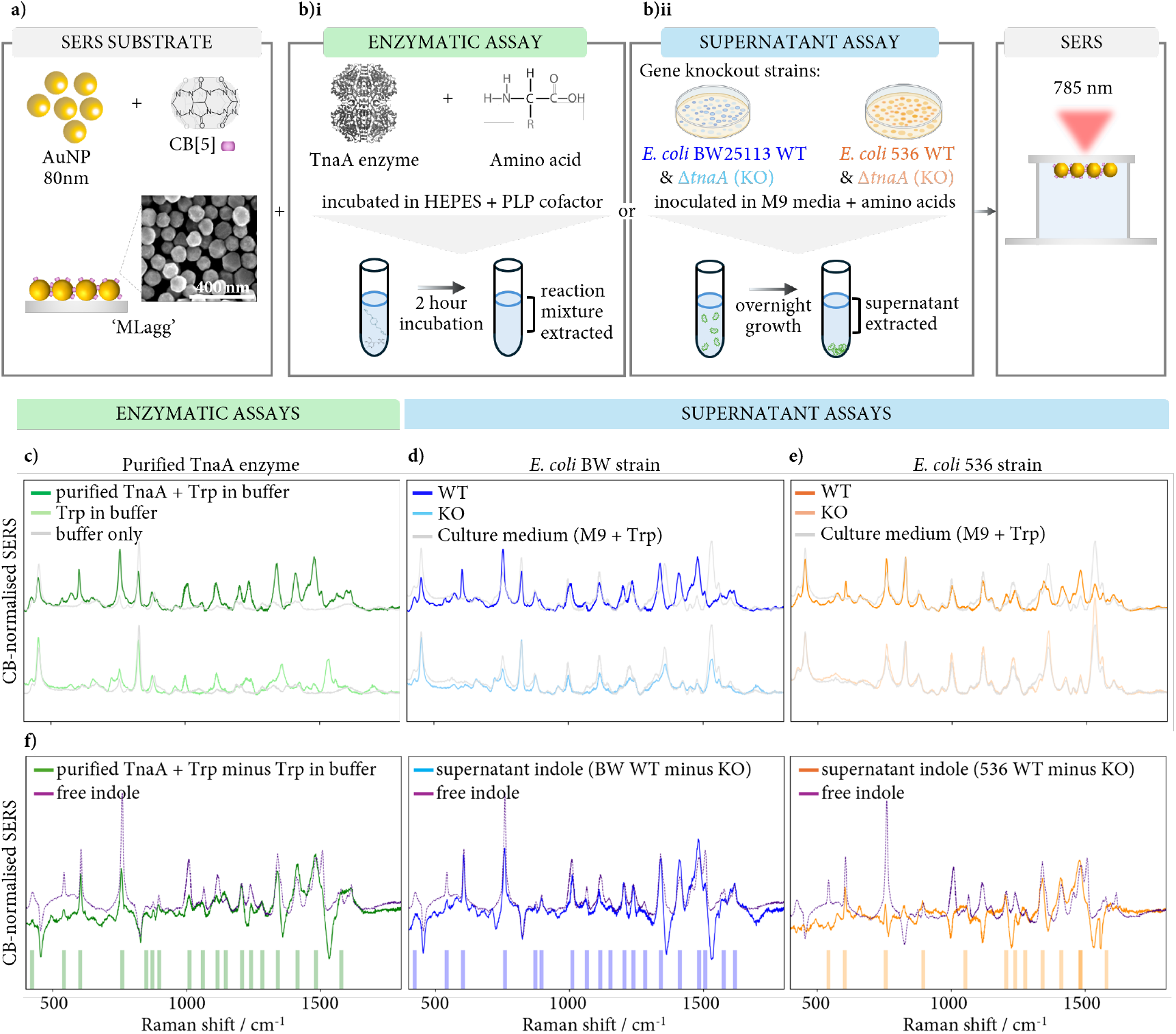
*In vitro* and *in vivo* SERS characterisation of TnaA-dependent indole production in *E. coli*. (a) SERS sensing films, from aggregation of 80nm gold nanoparticles with CB[5] molecules at a liquid-liquid interface before drying into films [43]. (b)i) Enzymatic assays which incubate the purified enzyme TnaA and amino acid in HEPES buffer and PLP cofactor; ii) *E. coli* grown overnight in media containing M9 media supplemented with amino acid (see Methods). Supernatant is extracted and SERS measured. (c-e) Normalised SERS spectra of (c) enzymatic assay of TnaA combined with tryptophan (dark green), an identical solution without the enzyme TnaA (light green), and buffer solution only (without tryptophan, gray); (d) *E. coli* BW25113 and (e) *E. coli* 536 of both wild-type (WT) and *tnaA* gene knockout strains (KO, Δ*tnaA*), together with culture media of M9 media supplemented with tryptophan shown for reference (gray). (f) Subtraction of the two spectra shown in (c-e) and comparison to the spectrum of 10mM free indole. Significant peaks shown as shaded vertical lines, demonstrating strong alignment between the spectra and presence of supernatant indole in these assays, as confirmed by mass spectrometry (see Fig. 4).

The goal of our present work is three-fold. First, we demonstrate how SERS with MLagg substrates can be used in combination with appropriately genetically-engineered strains for the quantitative, label-free, detection of indole within the high chemical complexity of a bacterial suspension, at nanomolar concentrations. Second, we show how to extend this methodology to investigate the broader cellular-based, TnaA-dependent metabolic activity for each of the 20 canonical L-amino acids, in order to provide mechanistic insights into bacterial physiology. Third, we leverage isotopomer-labelled substrates, such as tryptophan-d5, tryptophan-15N, indole-d6, together with amino acid isotopomers tyrosine-d4, leucine-d10, and valine-d8, to perform label-free structural deduction of an uncharacterised TnaA-dependent metabolite directly from cellular SERS spectra, without isolation or purification.

Our results demonstrate that applying SERS to target wild-type and knock-out strains facilitates the detection and quantification of even very low concentrations of byproducts of an enzymatic activity of interest, with minimal sample preparation and strong reproducibility. Isotope-resolved SERS extends this methodology from detection to label-free structural elucidation directly in living cell cultures. More broadly, our findings endorse SERS as the method of choice for sensitive, non-invasive, and inexpensive identification and quantification of metabolites in living cell cultures.

## Results

### Reconstruction of TnaA-dependent indole signature in *E. coli* cultures using SERS

Bacterial cell cultures are chemically complex, consisting not just of cells and cell debris, but also of all molecules from spent media and byproducts of bacterial growth. The isolation of a SERS spectrum for a target molecule in such complex solutions is challenging, as many compounds contribute with different vibrational modes.

To identify a clean SERS signature of indole in this environment, we perform SERS (Fig. 1) on the supernatant of two wild-type (WT) *E. coli* strains, BW25113 and 536, and their genetically modified counterparts, in which the *tnaA* gene has been knocked out (KO or Δ*tnaA*) (Fig. 1b ii), see Methods). Because the supernatants contain the spent media of the cell culture, these assays assess metabolite production from living cells. In parallel, we prepare an *in vitro* enzymatic assay comprising the purified enzyme TnaA incubated in a buffer containing tryptophan, and a corresponding control assay without TnaA (Fig. 1b i). SERS spectra for WT and KO supernatant assays for both *E. coli* BW25113 and *E. coli* 536 show strong peaks (Fig. 1d,e), compared to the culture media (shown in gray). Similar peaks are seen for the equivalent *in vitro* assay with and without TnaA (Fig. 1c). A clear difference in spectral profile is observed between WT and KO for supernatant assays, as well as with/without TnaA for enzymatic assays. This can be seen more clearly when looking at the spectral differences (Fig. 1f) in which the KO spectrum is subtracted from the WT spectrum to produce a clean TnaA-dependent supernatant spectrum (see Methods). The spectral profile of the difference spectra strongly matches the spectrum for free indole (10 mM, water, light crimson line in Fig. 1f), which is the known byproduct of tryptophan catalysis by TnaA (shared peaks highlighted by shaded vertical lines). The enzymatic assay exhibits an almost identical signature (Fig. 1f), demonstrating the ability of SERS to detect an excreted metabolite, such as indole, directly from a bacterial culture with minimal sample manipulation or extraction. The presence of indole in these supernatants is also confirmed independently by mass spectrometry (see Fig. 4). As expected, indole is produced from tryptophan by TnaA, and is not present when the enzyme TnaA is removed, as reflected in the SERS spectra. These results are further corroborated by spiking with 10mM of free indole in both strains (SI Fig. 1.1) and in artificial urine, a more complex media (SI Fig. 1.5).

Here, we reconstruct, for the first time, the indole spectrum using SERS directly in bacterial cultures without specific extraction [46, 47]. Importantly, this capability stems from comparing the wild-type strain with the appropriate corresponding genetically modified strain, where the gene (*tnaA*) encoding the enzyme of interest (TnaA) has been knocked out. This provides a control for the non-TnaA dependent metabolite activity. The second crucial factor is the size-selective filtering by the nanogap of only small molecules from the complex mixture, greatly improving the signal-to-noise ratio in the SERS spectra.

SERS from MLaggs can detect *in vivo* indole without specific extraction, at 100 nM concentrations (SI Fig. 1.4), a 500 times increase over prior work [46]. This holds significant promise for detecting indole at very low concentrations in clinical settings, particularly as indole is directly involved in over 20 infections, and the current limits of detection (3 *μ*M) using colorimetric assays are non-viable for low concentration detection [16, 48–50]. Calibration based on characteristic indole peak areas ((SI Fig. 1.4) shows a linear correlation (R^2^ = 0.93) between concentration and SERS intensities, from 100 nM to 10 mM.

### SERS assay profiling across amino acids reveals previously undetected TnaA-dependent activity

Having established that SERS can detect indole production directly from the supernatant of bacterial cultures and at very low (nM) metabolite concentrations, we extend this technique to all 20 canonical L-amino acids, to investigate potential metabolic activity of TnaA on other substrates. For enzymatic assays we isolate the effect of TnaA by subtracting the SERS spectrum of solutions of purified TnaA supplemented with the amino acid of interest, from the equivalent SERS spectrum without TnaA (*with TnaA* minus *without TnaA*, Fig. 2a). Similarly, for the supernatant assays, we compare the difference in SERS spectra from the supernatant of the WT cultures supplemented with the target amino acid, subtracting the SERS spectrum of the corresponding KO strain (WT minus KO, Fig. 2b,c). We perform this spectral analysis for both *E. coli* variants, BW25113 and 536. In summary, we utilise three types of assays (enzymatic, cellular *E. coli* BW25113 and cellular *E. coli* 536), two types of strains (WT, or with TnaA, and KO, or without TnaA), 20 amino acids, and ten repeats for each spectrum recorded, yielding a total of 1200 spectra (and a vector of size (1200, 4060)), which forms the basis for Fig. 2.

**Figure 2.**
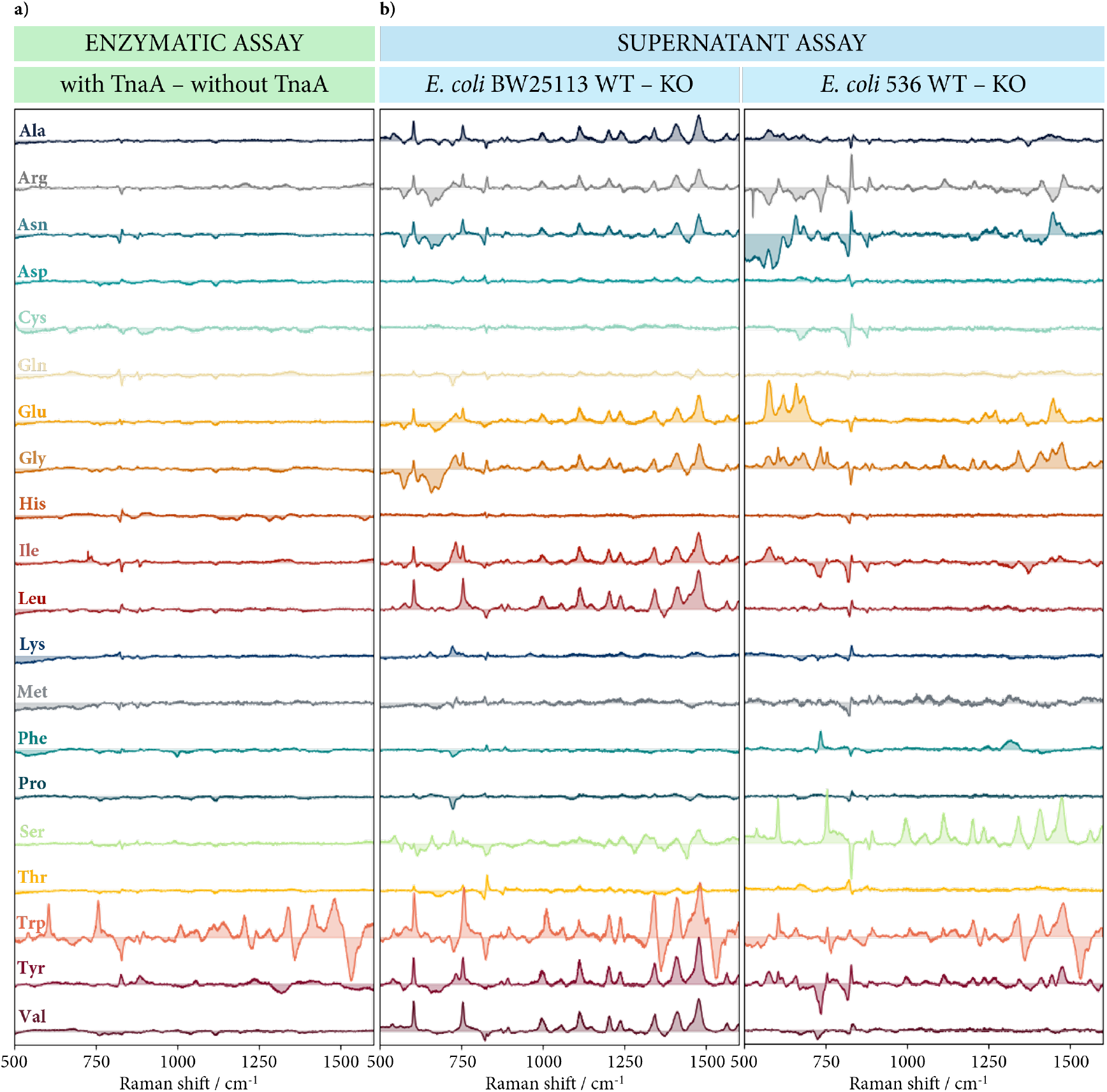
SERS profiling of enzymatic and supernatant cellular assays. (a) Enzymatic assays: SERS profiling with and without TnaA. (b,c) Supernatant assays: wild-type (WT) minus *tnaA* gene knockout (KO) assays, inoculated from LB agar into M9 media supplemented with each amino acid, as labelled (see left of figure).

In the enzymatic assay, only tryptophan exhibits significant TnaA-dependent metabolic activity, implying that this is the only amino acid directly targeted by TnaA with a byproduct detectable by SERS. It is known that both serine and cysteine are also TnaA targets, however, the byproducts of these reactions fall outside the fingerprint region considered here (500-1750cm^−1^) [24]. Surprisingly, we also observe substantial TnaA-dependent metabolic activity for several other amino acids in the supernatant assays, suggesting a previously uncharacterised role for TnaA in multiple metabolic pathways within a living bacterial cell. Amino acids L-alanine, L-isoleucine, L-leucine and L-valine show SERS activity in *E. coli* BW25113 but not in *E. coli* 536 or enzymatic assays. Conversely, L-arginine, L-asparagine, L-glutamate, L-glycine, L-serine, and L-tyrosine show SERS activity in both strains. L-serine and L-cysteine, which are known substrates of TnaA, show higher activity in *E. coli* 536 than *E. coli* BW25113, and no activity in either, respectively. To our knowledge, this is the first time that the role of TnaA in *E. coli* cultures supplemented with each of the twenty canonical L-amino acids has been characterised.

### PCA identifies a shared indole-like metabolic signature across amino acids

To perform a quantitative comparison between the activity of TnaA on different amino acids, we apply PCA on the SERS spectra shown in Fig. 2 (see Methods and SI Fig. 3.1-3.4). Projection of the data on PC1 and PC2 (Fig. 3, enzymatic, supernatant *E. coli* BW25113 and supernatant *E. coli* 536) shows a surprising trend. By superimposing the projections of free indole (indigo stars) and pure tryptophan (orange star) spectra on this two-dimensional space, we see PC1 mostly captures the enrichment/depletion of indole, while PC2 captures the enrichment/depletion of tryptophan in the solution, as confirmed by PC1 and PC2 weightings (SI Fig. 3.5). The position of the tryptophan enzymatic assay (orange square datapoint in Fig. 3a) confirms its enrichment of indole and depletion of tryptophan, consistent with the previously characterised role of TnaA in transforming tryptophan into indole. As suggested by the SERS spectra in Fig. 2, no other amino acid shows spectral activity in enzymatic assays, confirming that tryptophan is the only direct substrate of TnaA detectable by SERS in this context.

**Figure 3.**
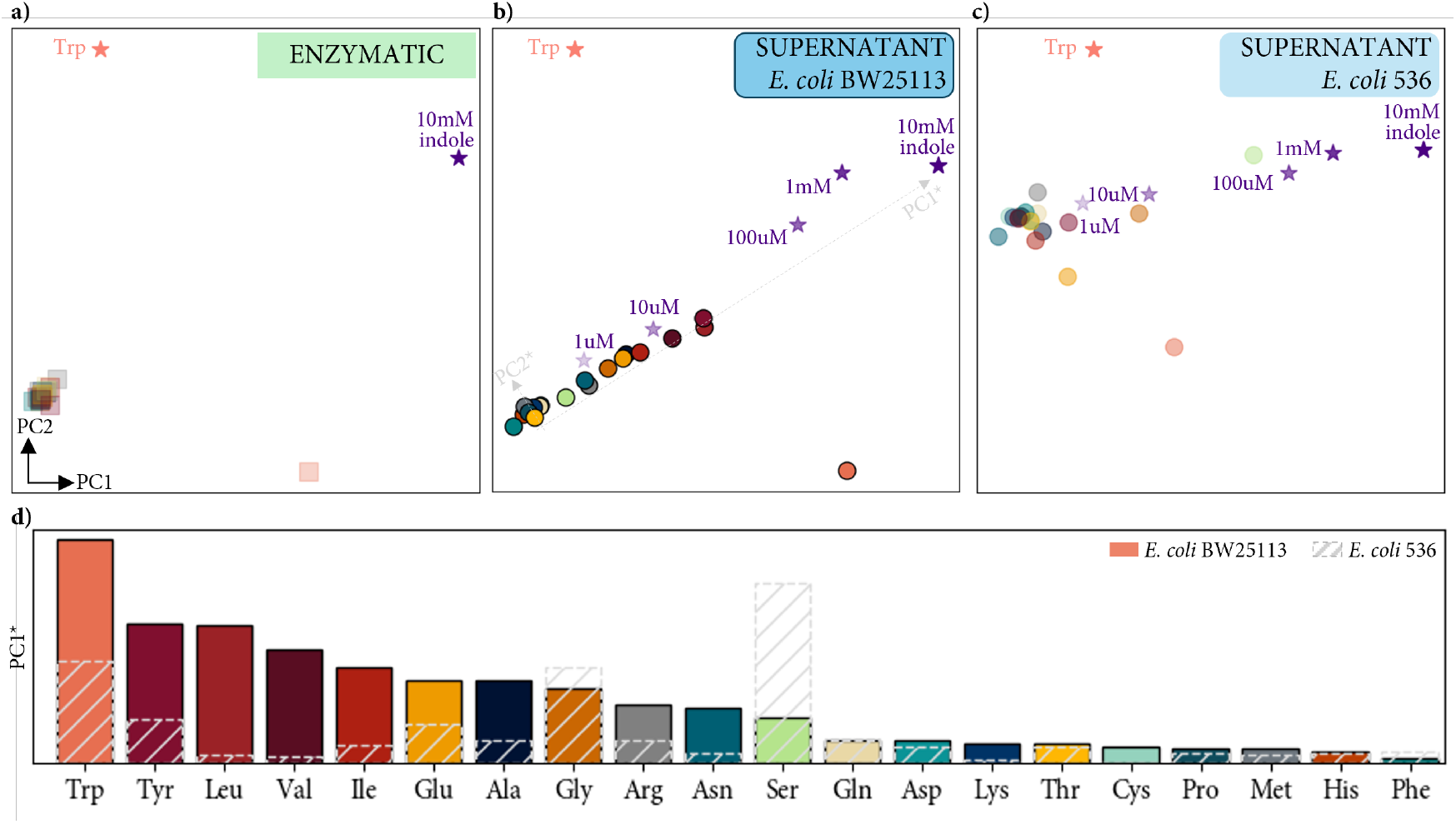
PCA of supernatant WT minus KO spectra. (a) PC1-PC2 projections of enzymatic assay spectra obtained by subtracting assay spectra without the enzyme TnaA from corresponding spectra with TnaA, and (b,c) of cellular wild-type (WT) spectra minus corresponding *tnaA* gene knockout (KO) assay spectra for (b) *E. coli* BW25113 and (c) *E. coli* 536, coloured by amino acid as in Fig. 2. (d) Ranking of each cellular amino acid assay, based on a rotated PC axis, which can be seen in gray lines in b). First two PCA components explain 65% of the variation (see Methods and SI Fig. 3.1-3.4).

The PCA for the two supernatant assays (Fig. 3b and Fig. 3c) shows a very different picture. First, tryptophan is similarly positioned across the three assays, confirming that the direct enzymatic reaction between TnaA and tryptophan is the main metabolic pathway when bacterial cells are exposed to tryptophan. However, all the other amino acids are found distributed along a line on the PC1-PC2 plane (see SI Fig. 3.1). The differences between enzymatic and supernatant assays clearly indicate that within the bacterial cell, there are indirect interactions between TnaA and amino acids beyond tryptophan. In addition, the observed linear trend raises the intriguing hypothesis that a common metabolite(s) might be excreted by the cell at different concentrations depending on the amino acid initially present in the media.

Given that the SERS spectra of WT minus KO assays for different amino acids predominantly extends along the free indole projection, we hypothesise the presence of a previously undetected indole-like metabolite, I*, in the supernatant of *E. coli* assays. To support this hypothesis, we project the SERS spectra of free indole at different concentrations onto the PC1–PC2 plane (indigo stars, Fig. 3a–c), which intriguingly fall along the same direction outlined by the amino acids. These results show that this distinctive indole-like signature is not unique to a particular *E. coli* strain or amino acid.

To quantify how similar the WT minus KO *E. coli* assay spectra are to the free indole spectrum, we rotate the PC1-PC2 projection so that the *x*-axis runs along the free indole direction (defined as PC1*, dashed line in Fig. 3d). We then determine the distance of each amino acid spectrum from the origin, here defined as the centroid of assays without an I* spectral signature (null region, Fig. 3d and Methods). Fig. 3d reports the PC1* values or horizontal distance from the null region in the rotated PCA. As expected, tryptophan gives the strongest signal, followed by serine for the *E. coli* 536 strain, and tyrosine and the BCAAs for the *E. coli* BW25113 strain. The other amino acids follow, although no obvious biochemical pattern can be determined in their ranking. Alternative measures of analysing difference spectra with different controls have also been explored (see SI Fig. 2.1-2.5).

### LC-MS of supernatant assays

To investigate the potential identity of I*, the indole-like signature detected by SERS across specific amino acid-supplemented cultures, we perform liquid chromatography–mass spectrometry (LC-MS) on selected supernatant assays for both WT and KO strains (see Methods). Fig. 4a shows the change between WT and KO as ΔMS = log_2_(WT/KO) of normalised peak area intensities for indole and its derivatives found in reference libraries. Only metabolites above the limit of detection (>3*σ*) are included. Unsurprisingly, for indole (top left chart, Fig. 4a), both *E. coli* BW25113 and *E. coli* 536 assays in tryptophan show significant ΔMS changes, while no meaningful differences are detected in the other WT vs. KO assays. The indole derivative 3-(2-hydroxyethyl)indole shows elevated levels in some assays that exhibit the indole-like signature (BW-tyrosine, 536-leucine, 536-glycine), but not in others (BW-leucine), and also appears elevated in assays with a very weak indole-like signature (BW-cysteine). Based on LC-MS, we find no single molecule (indole-related or otherwise) or combination of molecules displaying a common profile across the amino acids identified by SERS, or meaningful correlation with the I* spectrum observed (Fig. 2).

**Figure 4.**
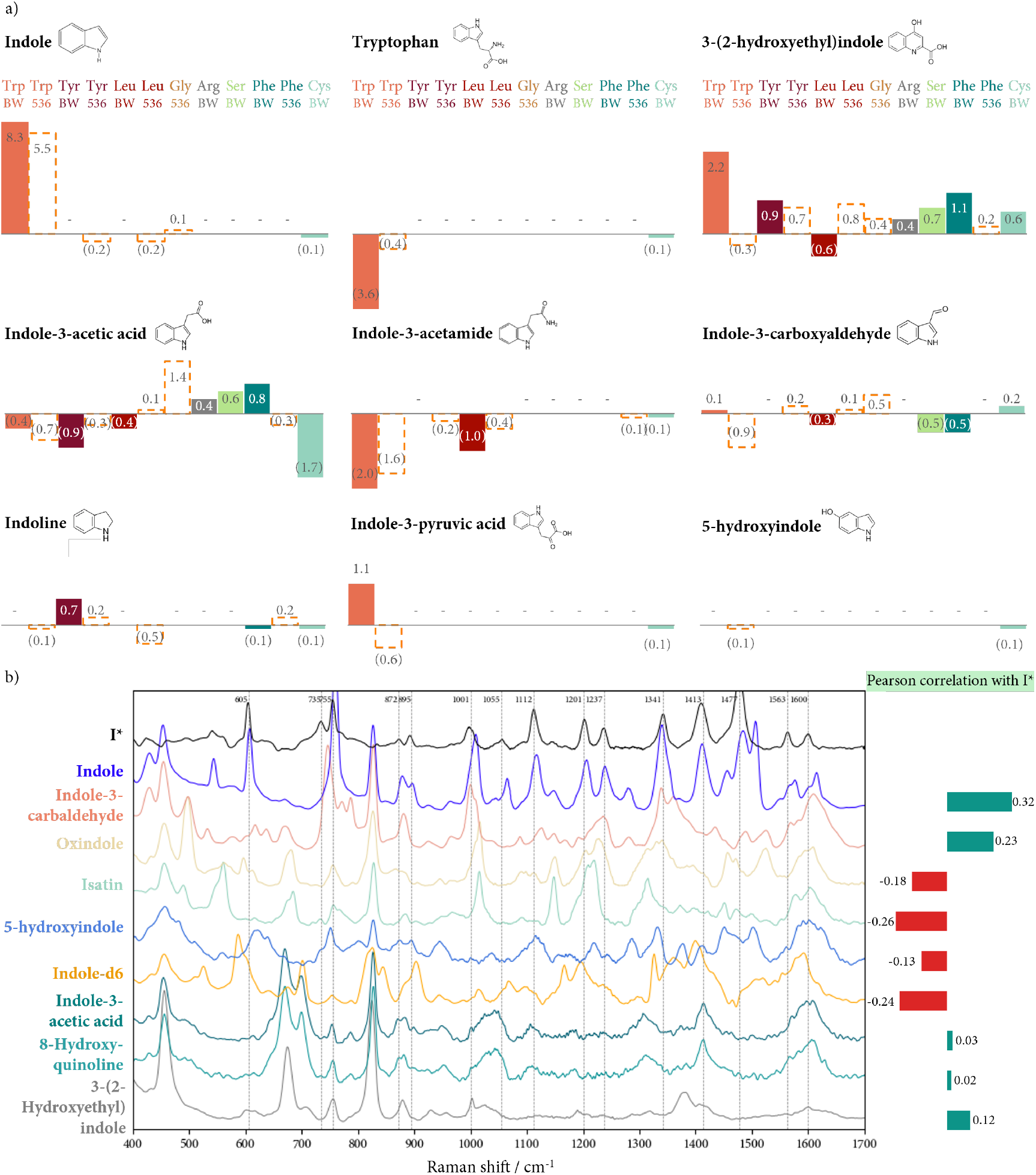
LC-MS and SERS results of indole derivatives. (a) LC-MS results (ΔMS = log_2_(WT/KO)) for the detection of indole and indole derivatives in selected supernatant WT minus KO cellular assays (*E. coli* 536 strain assays shown dashed). (b) Comparison of SERS spectra for candidate indole derivatives versus I* and indole. Pearson correlation highest with indole.

Overall, 914 metabolites are detected by LC-MS (see SI Fig. 4.1), with considerable variability, both across strains for the same amino acid (e.g. tryptophan), and across amino acids for the same strain (e.g. arginine and serine). Further analysis of the metabolite set in cultures supplemented with amino acids reveal potentially interesting shared metabolites. For example, the aromatic amino acids (tryptophan, tyrosine, phenylalanine) share trimethylamine N-oxide (TMAO), a clinically important metabolite found to play a role in cholesterol metabolism, and glycylproline, a metabolite released during the breakdown of collagen (see SI Fig. 4.2c)[51, 52].

Next, we perform SERS on nine candidate indole derivatives - oxindole, isatin, indole-3-acetic acid, indole-3-carbaldehyde, 3-(2-hydroxyethyl)indole, 5-hydroxyindole, 8-hydroxyquinoline, and the isotope-labelled reference indole-d6 - which overlap with the LC-MS indole panel identified. Fig. 4b compares the SERS spectra of I*, indole and these candidate derivatives, including the Pearson correlation with I*. No candidate yields a strong match: correlations range from +0.32 (indole) and +0.23 (indole-3-carbaldehyde), to -0.26 (isatin) and -0.24 (indole-d6). Together, the LC-MS and SERS results indicate that I* does not correspond to any of the common indole derivatives previously characterised. In fact, I* aligns more closely with the free indole spectrum. Relative to the spectrum for free indole, however, there are key differences, which we explore next (see Fig. 5 below). These results support our hypothesis that the previously unreported TnaA-mediated metabolic signature shared across several cultures supplemented with amino acids other than tryptophan is likely an indole derivative.

**Figure 5.**
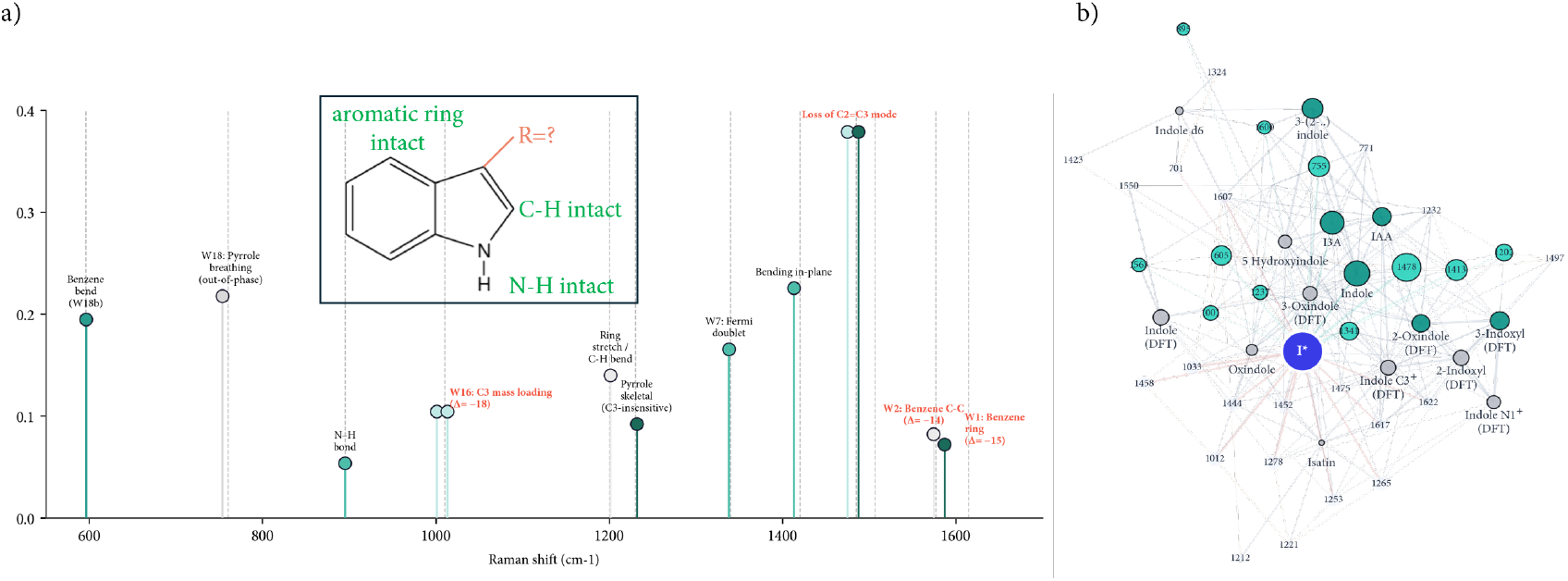
Peak comparison and SEM analysis predicts I* is consistent with a C3-substituted indole scaffold. (a) SERS band assignment of free indole (vertical gray lines) and I*, with line height reflecting relative intensity in I*. Band labels follow the W-mode nomenclature for indole derivatives (Dieng & Schelvis, 2010, Harada et al., 1986; Maruyama & Takeuchi, 1995). Orange bands highlight I*-specific diagnostic features: absence of C3-H in-plane mode at 1478 cm^−1^ is consistent with C3 substitution; redshift of Δ ≈ −18 *cm*^−1^ at W16 (1001 *cm*^−1^) indicate effective (non-H) mass at C3; and redshifts of Δ ≈ −13*cm*^−1^ at W2 (benzene C-C, 1563 *cm*^−1^) and W1 (benzene ring, 1600 *cm*^−1^) modes suggest electron-donating C3 substituent perturbing the benzene ring. Present bands confirmed at p ≤ 0.05 (Mann-Whitney U, for both BW25113 and 536 strains, n = 14 I*-positive spectra). (b) SEM-informed bipartite network representing structural relationships between I* (central node), SERS band positions (teal nodes, sized by I* intensity at that band), and indole derivative reference compounds (dark green nodes indicate positive correlation, grey nodes indicate negative correlation; nodes sized by SEM-inferred similarity to I*). Green edges indicate bands present in I*; red edges indicate absent bands. The network is constructed from L2-normalised band-molecule correlations weighted by SEM path coefficients (*γ*, see SI Fig. 5 for more details).

### Peak assignments and hypothesis of molecular structure of I*

To explore the molecular differences between indole and I* using SERS, we compare previously determined band assignments of free indole to I*, using the *E. coli* BW25113 tyrosine WT minus KO spectrum as best representing a clean I* signature. As Fig. 5a shows, several characteristic indole bands are preserved at their canonical positions in I*: the benzene C-C bending mode at 605 cm^−1^ (W18b), the out-of-phase pyrrole ring breathing mode at 757 cm^−1^ (W18), the N-H out-of-plane bending mode at 895 cm^−1^, and the ring stretching/C-H bending modes at 1201 and 1232 cm^−1^ [53]. The conservation of these modes indicates the bicyclic indole scaffold and the N-H bond are retained in I*, and that the aromatic C-H bonds at positions 2, 4, 5, 6, and 7 are intact.

In contrast, three classes of bands diverge from free indole in I*. First, the C3-sensitive ring breathing mode (W16) is red-shifted from 1019 cm^−1^ in free indole to 1001 cm^−1^ in I* (Δ = −18 cm^−1^), indicating effective mass loading at the C3 position of the ring [47, 53–55]. Second, the C2=C3 stretching mode at 1507 cm^−1^ and the C3-H in-plane bending mode at 1278 cm^−1^ are absent from I*, consistent with a C3 substituent [47, 54, 55]. Third, the benzene C-C stretching modes (W2 and W1, at 1577 and 1615 cm^−1^, respectively) are red-shifted in I* by −14 and −15 cm^−1^, respectively, reflecting electronic perturbation of the benzene ring by the C3 substituent [**walden**, 53]. We note however that these experimental spectra arise from molecules within the MLagg substrate trapped in nanogaps at Au facets, which is known to shift analyte peaks and may additionally depend on environmental factors (pH, solvation) that cannot yet be easily accommodated or controlled for in experiments. Taken together, these three signatures - mass loading at C3, loss of C3-H, and ring-mode perturbation consistent with a conjugated electron-donating substituent - point to I* comprising an indole scaffold with a substituent at the C3 position (Fig. 5a, inset). These assignments are further evaluated by isotope-resolved analysis (Fig. 6 below).

**Figure 6.**
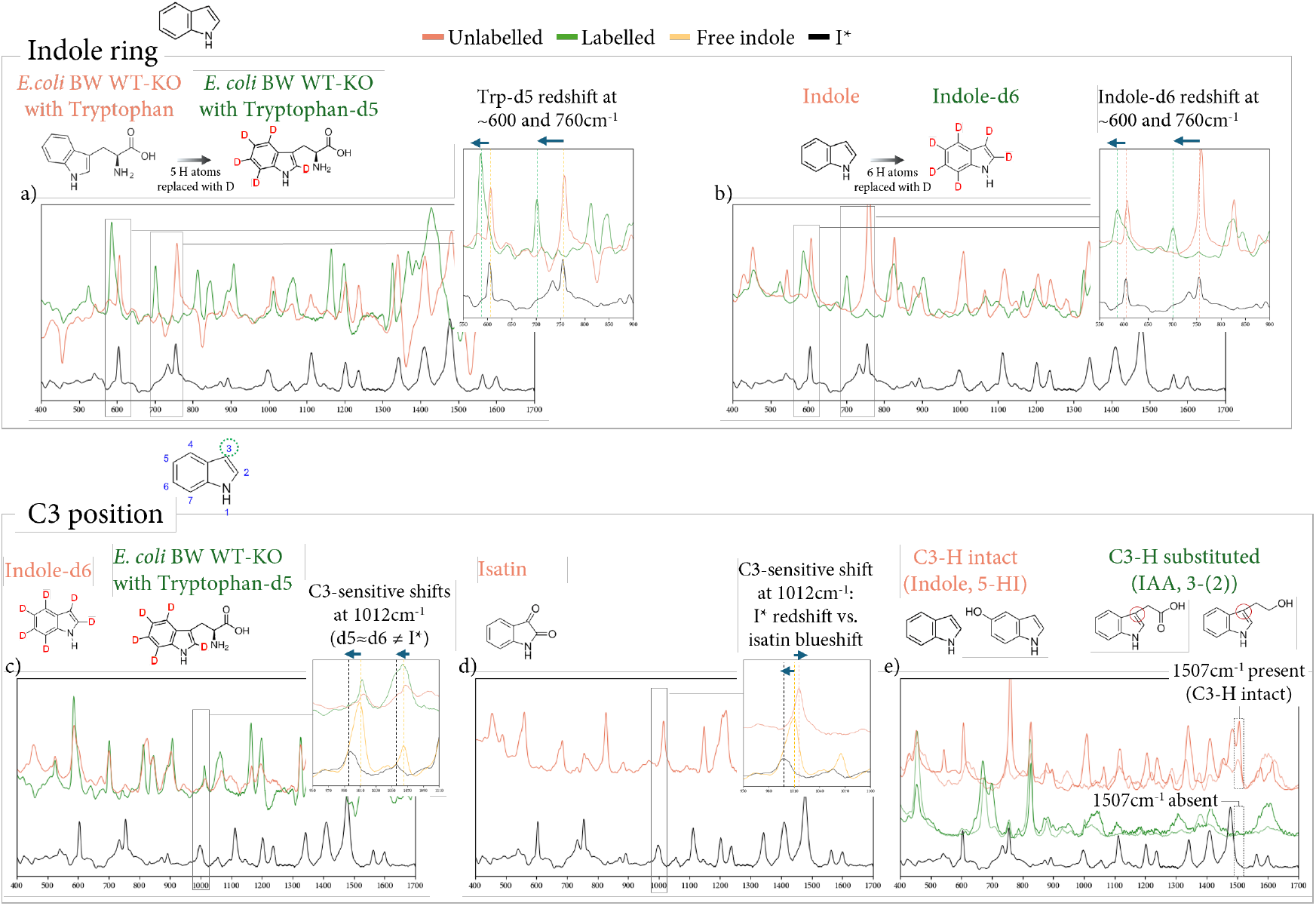
Isotope and structural reference controls point to I* comprising an intact indole ring with modification to the C3 position. (Top panel) Comparison of two characteristic peaks of indole at 605 cm^−1^ (benzene C–C bending mode) and 760 cm^−1^ (out-of-phase pyrrole ring breathing mode) showing the bicyclic indole scaffold is preserved in I*. (a) Comparison of *E. coli* BW25113 WT minus KO supernatant spectra cultured in tryptophan versus tryptophan-d5, showing d5-induced red-shifts of these peaks. (b) Corresponding shifts in indole-d6 relative to free indole and I* confirm these peak assignments and demonstrate I* likely retains aromatic C–H bonds at these ring positions, like free indole. (Bottom panel) Evidence that the I* modification is localised to C3. (Left) C3-sensitive modes at 1012 cm^−1^, where I* is red-shifted and where indole ≈ d5 ≈ d6, indicating mass loading at C3. b) Comparison with isatin showing the same 1012 cm^−1^ mode blue-shifted in isatin, but red-shifted in I*. (e) C2=C3 stretching mode at 1507 cm^−1^ absent from I*, consistent with other C3-H substituted indole derivatives such as IAA and 3-(2-hydroxyethyl)indole. Boxes and dashed lines mark diagnostic spectral regions.

### Isotopes and structural references elucidate structure of I*

We next deploy a series of isotope and structural reference controls to validate the proposed structure of I* (see Fig. 5). Fig. 6 shows the results. By comparing the SERS spectra of *E. coli* BW25113 WT minus KO supernatants cultured in tryptophan versus tryptophan-d5 (Fig. 6a), we observe substantial red-shifts of the benzene C-C bending mode at 605 and the out-of-phase pyrrole ring breathing mode at 760 cm^−1^ in the tryptophan-d5 difference spectrum, with corresponding shifts between free indole and indole-d6 (Fig. 6b). Because indole-d5 comprises labelling of deuterium at ring positions 2, 4, 5, 6, and 7, and not C3, via TnaA-mediated cleavage of tryptophan-d5, these shifts establish that the aromatic C-H bonds at non-C3 positions are intact in I*.

Fig. 6 c), d) and e) provides evidence the modification to the indole ring is localised at C3. First, as shown in Fig. 6c, the C3-sensitive ring breathing mode (W16) at 1012 cm^−1^ is red-shifted in I* to 1000 cm^−1^, while free indole, indole-d5, and indole-d6 all cluster together at 1012 cm^−1^. Because indole-d5 and indole-d6 differ only at the C3 site (where indole-d6 is labelled and indole-d5 is not), the agreement of both peaks establishes that this mode is insensitive to C3-H bending. The shift in I* therefore cannot arise from altered C3-H character and instead may reflect mass loading by a non-hydrogen substituent at C3.

Fig. 6d shows the comparison with isatin and reveals that the same 1012 cm^−1^ breathing mode is *blue*-shifted in isatin (likely due to ring stiffening from the ring-internal C3=O) but *red* -shifted in I*. The contrasting shifts demonstrate the C3 modification in I* may not be a ring-internal oxidation, as in isatin, but an exocyclic substituent attached to C3. Finally, the C2=C3 stretching mode at 1507 cm^−1^ is absent from I* (Fig. 6e) [54]. This mode is sensitive to C3 substitution; in free indole this mode involves displacement of both C2 and C3 atoms [54]. Thus a C3 substituent may likely perturb this mode through mass loading at C3 as well as altered C2=C3 bond order. This is also evidenced by the presence of this peak shifted upwards to 1552 cm^−1^ in tryptophan, where C3 carries an exocyclic carbon rather than H [54, 55]. Its absence from the 1500 − 1530 cm^−1^ region in I* therefore directly reports on a perturbed C2=C3 bond environment, consistent with C3 substitution.

### Unaltered I* signature with amino acid isotopomers suggests indirect I* production pathway

Building on the structural insights shown in Fig. 6, we determine whether the amino acids producing the strongest I* signatures contribute directly to its molecular structure, and are being cleaved by the TnaA enzyme. We repeat the *E. coli* BW25113 supernatant assays, this time using isotopomer-labelled tyrosine, leucine and valine, comparing tyrosine/tyrosine-d4, leucine/leucine-d10 and valine/valine-d8 supplementation (Fig. 7). Across all three amino acid pairs, the labelled and unlabelled spectra remain essentially unchanged at the diagnostic I* bands, including those at 605, 760, 1012 and 1480 cm^−1^. This contrasts with the clear isotope-dependent shifts observed for tryptophan-d5 (upper-most spectrum, Fig. 7) and indicates that the I* carbon skeleton does not derive directly from tyrosine, leucine or valine. Instead, these amino acids appear to promote I* formation indirectly, most likely by reallocating intracellular amino acid metabolism and increasing availability or processing of an intracellular tryptophan-derived pool through TnaA.

**Figure 7.**
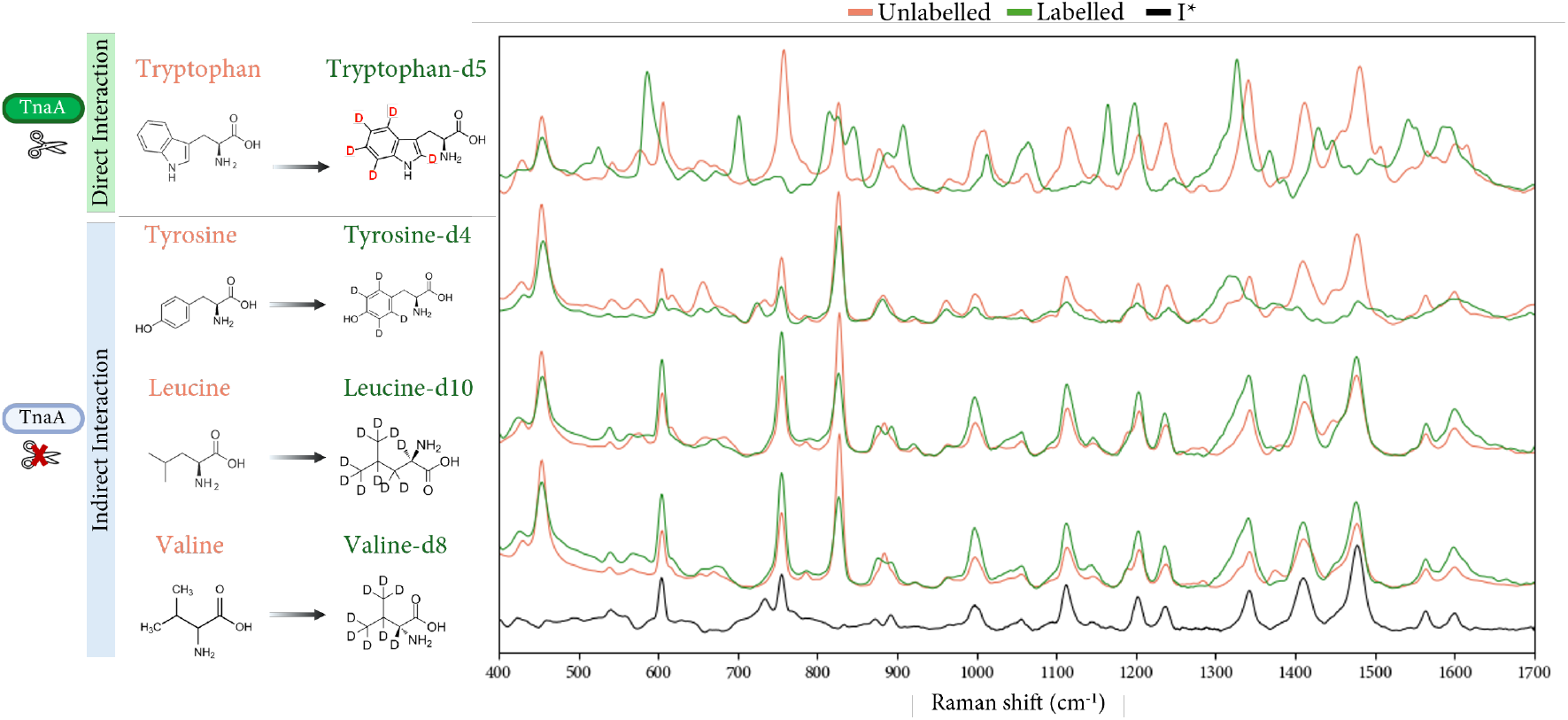
Isotope-labelling controls for amino acids producing the largest I* signal (tyrosine, leucine, and valine) confirm that I* is likely derived from intracellular tryptophan. SERS spectra of E. coli BW25113 WT supernatant cultured in M9 medium supplemented with each unlabelled amino acid (red) or its deuterated equivalent (green). I* reference spectrum (black) shown for comparison. Across all three amino acid pairs, the labelled and unlabelled spectra are not meaningfully different at band positions attributable to I*, which include indole ring modes at 605, 760, 1012, and 1480cm^−1^ that shift under tryptophan-d5 labelling (see Figure 5). There is no meaningful displacement in I* at each amino acid’s characteristic peaks, indicating the carbon skeleton of I* contains no atoms originating from these amino acids. This points to the origin of I* to intracellular tryptophan, and excluding contributions from aromatic (tyrosine) or branched-chain (leucine, valine) amino acid metabolism.

We propose two potential production mechanisms for I*, arising either from the TnaA-mediated metabolism of a derivatised form of tryptophan (3a in Fig. 8), or the TnaA-mediated production of I* via intracellular indole modification (3b in Fig. 8). Given the tight regulation of intracellular tryptophan levels, we speculate that supplementation with certain amino acids may perturb its balance, leading cells to reallocate metabolic flux toward utilisation of the externally supplemented amino acid, thus increasing relative intracellular tryptophan concentrations [56]. Fig. 8 illustrates a mechanistic example with leucine (which generates the second largest I* signal after tyrosine), based on known biosynthesis pathways [57]. The leucine-responsive regulatory protein (Lrp) is a key regulator of amino acid metabolism in *E. coli*. Lrp binding with leucine changes its conformation, binding differently to the promoters of Lrp-regulated operons [57]. This indirectly or directly downregulates *sdaA, sdaB, tdh* and other genes involved in the metabolism of other intracellular amino acids such as serine and threonine, resulting in a lower incentive for the cell to harvest aromatic amino acids such as tryptophan for nitrogen supply, thus increasing relative intracellular tryptophan concentrations [58]. More broadly, we speculate that an increase in branched chain amino acids (BCAAs) supplementation may shift the metabolic balance away from tryptophan breakdown for nitrogen, as the cell reoptimises use of leucine and other BCAAs to produce glutamic acid as its main nitrogen source. The schematic in Fig. 8 also shows the likely concentration of I* in the outer membrane rather than in solution. Free indole has been shown to have a 90-fold higher affinity for lipids, due to hydrophobicity of the aromatic ring [59]. Thus we posit that most of the I* metabolites could be potentially absorbed by cells (discarded in our assay), leaving low (nanomolar) residual concentrations of indole derivatives in the supernatant, which SERS can detect but existing methods cannot (see Discussion).

**Figure 8.**
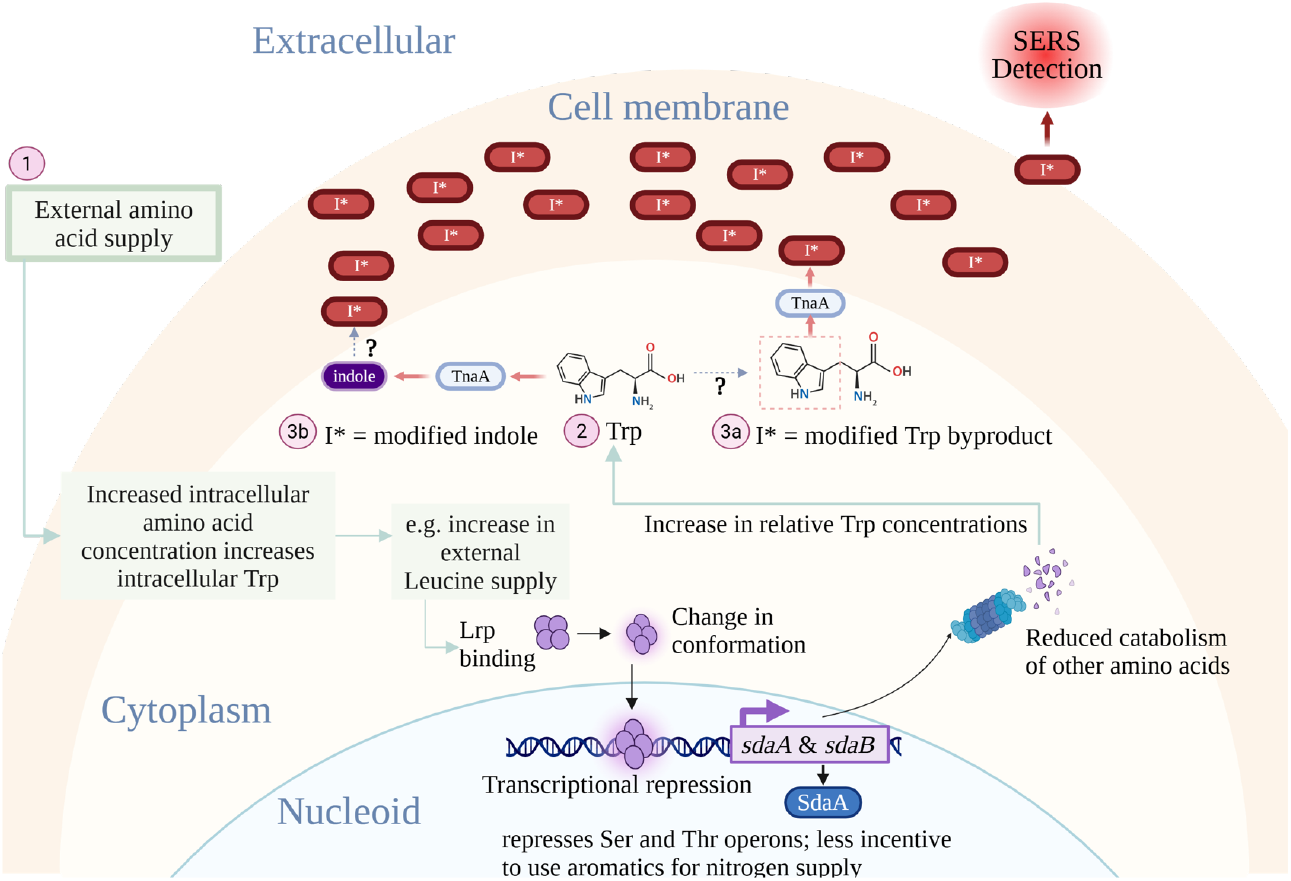
Schematic illustrating potential production mechanism for I* and role of amino acids. External supplementation of amino acids perturbs metabolic flux, leading cells to reallocate intracellular levels of amino acids, including tryptophan (see pathway labelled 1). Increased relative tryptophan concentrations activates TnaA, which leads to the production of I* either via a derivatised tryptophan (3a), or a modified intracellular indole (3b). Higher affinity of indole derivatives for lipids may leave majority of I* in cell membrane (which we discard), and low residual supernatant concentrations.

## Discussion

Our results support the finding that a previously unreported TnaA-mediated metabolic signature (I*) shared across several amino acids is an indole-like derivative with spectral features consistent with C3-localised modification, produced either from intracellular indole or from a derivatised form of tryptophan. We speculate that its presence has so far been undetected due to 1) the current limits of detection of both LC-MS and colorimetric biochemical assays relative to nanoplasmonic SERS, and 2) the 90-fold higher affinity of indole and indole derivatives for lipids in the cell membrane versus solution, resulting in low extracellular concentrations of I*.

In this work, we have demonstrated a SERS-based methodology for the *in vivo*, ultrasensitive detection of indole, and discovered an indole derivative in multiple assay types, from enzymatic to cellular assays, using SERS when extending this methodology to all 20 canonical L-amino acids. These results contribute to efforts in mapping previously overlooked bacterial signalling pathways which can have importance for clinical diagnostics [48, 49]. Recent work by Linares-Otoya *et al*., for example, characterised the N-acyl-cyclolysine (ACL) system in bacteria using a combination of techniques including LC-MS, pointing to potentially important insights into microbiome-host interactions [60].

I* has a peak profile which strongly points to an indole derivative, with an intact bicyclic ring scaffold and a C3 substituent (see Fig. 5-7). The presence of I* was further validated by performing replicate experiments with additional cell-washing steps (washed three times in PBS prior to inoculation). This series of experiments was performed to exclude the possibility that any indole transferred from cells from LB agar plates on which cells were initially streaked and grown was responsible for the observed signal. We find minimal spectral differences between washed and unwashed assays (see SI Fig. 2.8). Additionally, the abundance of I* is not consistent across cellular supernatant assays supplemented with amino acids, which reinforces our conclusion that specific amino acids may modulate the formation, degradation, or utilisation of indole or indole-like derivatives.

Furthermore, to verify that I* is not simply indole at very low concentrations such that the limit of detection of the colorimetric indole assay (3*μ*M) prevents its detection, or indeed an artefact caused by SERS matrix effects, where free indole may be oxidised by hot carriers generated from the excitation of surface plasmons, we concentrated hydrophobic metabolites in the cellular supernatant by >50x using C18 solid phase extraction cartridges, thus pushing the limit of colorimetric assay-based detection to 0.1 *μ*M (see Methods). No indole was detected in these colorimetric assays using a modified version of Kovács reagent (QuantiChrom™ Indole Assay Kit, BioAssay Systems). Kovács reagent contains p-dimethylaminobenzaldehyde (DMAB), which binds at the 3-position of the pyrrole ring, suggesting any indole derivative present is modified in this region [61]. Furthermore, the LC-MS results demonstrate no match between I* and common indole derivatives.

Some work has shown that tryptophan derivatives can be converted into oxidised indoles [62, 63]. Such oxidative products of tryptophan metabolism have important biological effects on bacterial signalling and behaviour, including the activation of signalling pathways that regulate immune cell differentiation and function via transcription factors such as the aryl hydrocarbon receptor (AhR), or protective roles in cardiovascular and metabolic diseases [62, 64].

The isotope-resolved experiments shown in Figs. 5-7 refine this view and show that I* retains an intact bicyclic indole scaffold, while the behaviour of C3-sensitive modes, the opposite shift direction relative to isatin, and the loss or perturbation of the C2=C3 stretching mode, point to modification localised around the C3 position rather than ring-internal oxidation. This distinction is important because it suggests that I* is not simply an oxidative degradation product of indole, but potentially a structurally specific TnaA-dependent indole derivative whose formation may involve an exocyclic C3 substituent. In parallel, the absence of any meaningful displacement of I*-associated bands in tyrosine-d4, leucine-d10 or valine-d8 supplementation experiments argues against direct incorporation of atoms from these amino acids into the I* carbon skeleton. Taken together, these isotope results support a model in which amino acid supplementation alters intracellular metabolic allocation, thereby modulating the availability or processing of a tryptophan-derived pool that is converted through TnaA-dependent activity into I*. More broadly, this highlights the value of SERS not only as a sensitive detection method, but as a structural probe capable of testing biosynthetic origin directly in complex bacterial supernatants without prior isolation of the metabolite.

The results from our enzymatic assays demonstrate that within the spectral range detectable by SERS, only tryptophan is identified as a direct substrate of TnaA (although serine and cysteine are also known substrates). Our cellular assays, however, report TnaA-mediated production of I* when the bacterial culture is supplemented with amino acids beyond tryptophan. Because the enzymatic assays rule out a direct interaction between these amino acids and TnaA, combined with the results of our amino acid isotope experiments (Fig. 6), we conclude that the effect of these amino acids on the activity of TnaA must be indirect. Given the tight control on the internal tryptophan concentration, we speculate that the external supplementation of particular amino acids either alters the internal pool of tryptophan, creating an excess of tryptophan and tryptophan derivatives that TnaA converts to indole, or activates TnaA’s expression triggering tryptophan conversion even at low tryptophan concentrations, and results in intracellular indole which is modified to I*. The second option seems unlikely given that it has been demonstrated that constitutively expressing high levels of *tnaA* does not result in significant degradation of internal pools of tryptophan [65]. In contrast, amino acid syntheses share multiple pathways and precursors, facilitating indirect interactions between the presence of one amino acid on the synthesis or availability of another [56]. For instance, tyrosine, which exhibits among the strongest I* signals in our results, can relieve repression of tryptophan synthesis by acting as an alternative substrate, indirectly promoting tryptophan production [66]. Similarly, serine is a precursor of tryptophan, thus its availability can increase tryptophan synthesis [67]. The supply of leucine, which displays a strong I* signal, may shift the metabolic balance away from energy-dense tryptophan degradation to produce glutamic acid for nitrogen (see Fig. 8).

Interestingly, we find that the positions of the amino acids within the biosynthesis pathways that result in indole derivative production reveal no clear or consistent pattern [56]. This suggests distinct regulatory mechanisms may underlie I* production for different amino acids. For example, isoleucine exhibits a strong I* signal, despite being synthesised downstream of threonine, which shows a minimal I* signal in SERS. Likewise, tyrosine displays a robust I* signal, whereas phenylalanine does not. Further investigation of these amino acid subgroups may provide valuable insights into the cellular strategies governing amino acid allocation. It is important to note that the concentration of I* released in the supernatant in these conditions is low, as expected given the 90-fold higher affinity of indole with lipids in the cell membrane versus solution, underscoring the key role that can be played by ultra-sensitive detection methods such as SERS. For instance, our work highlights how these methods can be used to investigate how the cell regulates and adapts its internal amino acid pool depending on the external conditions [59]. The differences in I* production between the two *E. coli* strains for the same supplemented amino acid also show that the optimal amino acid allocation that the cell maintains strongly depends on the genetic background, even for bacteria coming from the same species.

Finally, our findings on I* production in these unexpected conditions raise important questions on the role of indole and its derivatives as a signalling molecule among bacteria: how much of the I* production that we detect is simply a byproduct of optimising tryptophan cellular concentration versus being functionally employed for bacterial communication? Can other bacteria detect the level of I* produced in these conditions and respond to it? Can we associate or relate some of these conditions with the scenarios in which indole production is known to be important, such as biofilm formation or infections? Follow up studies in this direction have the potential to shed new light on the interplay between bacterial metabolism and ecology.

Previous characterisation of TnaA substrates (tryptophan, serine, and cysteine) has primarily been achieved through the use of enzymatic assays [24]. These assays rely on known reaction products as a readout of activity, which limits their utility in screening for interactions that result in unknown metabolic byproducts.

In contrast, LC-MS represents the gold standard for metabolomic studies investigating biochemical reactions by identifying their byproducts [68]. Despite its broad utility and high-throughput, its ability to identify unknown metabolites is restricted by existing annotated libraries [69]. In addition, LC-MS requires invasive sample manipulation and significant processing costs which often make it impractical for clinical applications[70].

Our work shows that combining SERS with targeted synthetic biology using clean gene knockouts bridges the gap between these two sets of techniques. On one hand, we show that SERS enables the ultrasensitive detection of previously uncharacterised molecular interactions, with very little sample preparation. In particular, SERS is found to be equally capable of detecting direct enzymatic activity *in vitro*, and enzyme-mediated activity *in vivo* with far superior sensitivity to other existing techniques. We achieve nanomolar-level sensitivity without specifically optimising for the detection limit, demonstrating the potential of current-generation SERS as a readily accessible approach for detecting indole in over 20 associated infections [48].

On the other hand, SERS provides a vibrational ‘fingerprint’ (unlike readouts from conventional *in vitro* assays) for metabolic byproducts of enzymatic interactions, which can be compared to existing libraries or tested against potential candidates to identify the possible molecule of interest, as is the case here. Importantly, even if band assignments are unclear and reaction byproducts cannot be identified, the precision of SERS can be used for comparative studies across different substrates (i.e. amino acids) or strains (i.e. WT vs. KO) to more accurately assess whether similar or different reactions may be at play across assays.

Crucially, we find that the comparison of WT and KO SERS spectra (Fig. 2), where the gene encoding the enzyme of interest (in this case TnaA) is knocked out, is key in isolating a clean SERS spectrum for assessing the enzymatic activity of a protein of interest within the complex chemical environment of a bacterial culture. Constructing gene knock-outs in most laboratory strains is relatively straightforward with several such libraries available, making this approach generalisable to many different enzymes or proteins of interest across multiple bacterial species. This work also opens new research possibilities and quantitative studies on the interplay between environment, bacterial physiology and specific biochemical reactions within the cell, orthogonal to current metabolomics efforts. For example, further applications to other gene-related modifications (e.g. knock-downs, integration of other genes) and/or to bacteria grown in different physiological conditions (e.g. nutrient concentrations, temperature, antibiotic susceptibility), combined with the ultrasensitivity of SERS has the potential to yield novel microbiological and clinically relevant insights. In particular, the SERS methodology outlined in this work may be particularly reliable and accessible for low density bacterial samples (e.g. environmental samples or animal/human microbiomes), where the metabolite(s) of interest might be present at very low concentrations. Additionally, SERS may help to shed light on more general real-time aspects of bacterial biochemistry. In a clinical context, the use of precision SERS may hold promise for the non-invasive, *in vivo* metabolite characterisation at the point-of-care, in a precise, reproducible and low-cost manner, positioning it as a potentially revolutionary tool in microbial diagnostics and public health.

## Materials & Methods

### Bacterial culture

*E. coli* BW25113 and *E. coli* 536 strains were used for all experiments. BW25113 Δ*tnaA* was obtained from the Keio collection [71]. 536 Δ*tnaA* was generated by *λ*-red recombination [72].

Bacterial freezer stock held at -80 °C was used to streak an LB agar-based plate, incubated overnight at 37 °C. Single isogenic colonies were collected with sterile plastic loops and used to inoculate 3 mL culture in M9 supplemented with 140 *μ*L of 1 mM amino acid (filter sterilised). This was incubated overnight at 37 °C. 500 *μ*L of the solution was extracted and centrifuged for 10 min at 7500g. The supernatant was extracted and SERS performed. Four biological replicates were prepared for all cellular assays. Of these, two biological replicates were derived from two single colonies taken from two different streak plates.

### Culture medium preparation

The M9 medium was prepared using autoclaved DifcoTM M9 Minimal Salts 5x, filter sterilised 20% glucose solution, 1M MgSO_4_ and 1M CaCl_2_. 14.1g of powder was dissolved into 250mL of sterile purified water (Milli-Q). The solution was autoclaved at 121 °C for 15 minutes. The autoclaved 5x solution was then diluted to 1x by mixing 50 mL into a 200mL volume of Milli-Q. 5 mL of 20% glucose solution, 0.5mL of sterile 1.0 M MgSO4 solution and 0.025 mL of sterile 1.0 M CaCl_2_ solution was added. 50mM of stock concentration of amino acids L-Cysteine (121.16g/mol), L-Arginine (174.2g/mol), L-Alanine (89.09g/mol), L-Asparagine (132.12g/- mol), L-Glutamic Acid (147.13g/mol), L-Glycine (75.07g/mol), L-Isoleucine (131.17g/mol), L-Methionine (149.21g/mol), L-Tyrosine (181.19g/mol), L-Tryptophan (204.23g/mol) and L-Serine (105.09g/mol) were prepared. For 3 mL of M9 medium, 140 *μ*L of each amino acid was added to give a 1mM final concentration.

Where specified, cells were cultured in Luria-Bertani (LB) broth (0.5 g/L NaCl), artificial urine, or grown on LB agar plates either containing no antibiotic or 50 *μ*g/mL kanamycin. Artificial urine media was prepared as described in supplementary methods.

### *λ*-Red recombination

*λ*-Red recombination was carried out to engineer the *tnaA* knock-out of 536 using the pSLTS helper plasmid [73]. Electrocompetent cells with arabinose-induced pSLTS were prepared as described in [73]. The kanamycin resistance cassette within *E. coli* BW25113 Δ*tnaA*:*kanR* was amplified by PCR, using primers designed to incorporate 91 bp and 96 bp homology arms of the sequences upstream and downstream of *tnaA* in *E. coli* 536 [**methods_5**]. This PCR product was purified using QIAquick® PCR Purification kit, then 100 ng was added to 50 *μ*L of the prepared electrocompetent cells, which were electroporated, recovered with 450 *μ*L of LB, then incubated overnight at 30 °C with 150 rpm shaking in a 15 mL tube. The following day, the cultures were centrifuged at 13,000 rpm for 1 minute, resuspended in 100 *μ*L of fresh LB, plated on LB agar containing 50 *μ*g/mL kanamycin, and incubated overnight at 37 °C without shaking.

The following day, single colonies from the transformation plate were screened for correct knockouts of *tnaA* by inoculating the single colonies in LB, incubating overnight at 37 °C with shaking, then using Kovac’s reagent to confirm the absence of indole production [74]. Following the confirmation of the absence of indole production, the strain was sent for whole genome sequencing.

### Enzymatic assay

Enzymatic assays were performed to assess direct TnaA-mediated degradation of individual amino acids. The highly active stock of TnaA (see supplementary methods for protein purification and preparation) in 0.1M HEPES (pH 7.8), 10% glycerol was concentrated using VivaSpin 500 centrifugal concentrators (10 kDa MWCO), then added to 500 *μ*L of 0.1 M HEPES (pH 7.8) containing PLP and 1 mM final concentration of a single amino acid, and incubated for 2 hours at 37 °C. For L-tryptophan, 80 *μ*g of TnaA and 0.5 *μ*M PLP were included. For L-cys, 400 *μ*g of TnaA and 20 *μ*M PLP were included. For L-ser and other L-amino acids, 400 *μ*g of TnaA and 5 *μ*M PLP were included. These amounts of TnaA and PLP were identified as sufficient for complete degradation of 1 mM L-trp, L-cys, and L-ser, within 1.5 hours at 37 °C using a lactate dehydrogenase-coupled enzymatic assay [75].

For controls, parallel reactions were set up as above, but adding the same volume of concentrator flow-through (0.1 M HEPES (pH 7.8), 10% glycerol) instead of TnaA. Enzymatic reactions were not filtered prior to SERS measurements, as tests comparing filtered reactions to unfiltered reactions demonstrated that purified TnaA provided negligible SERS signal relative to the buffer solution (see SI Fig. 1.6).

### Indole concentration and quantification

Indole in bacterial supernatant samples was concentrated and quantified using a modified version of the approach described by Zarkan et al [76]. Briefly, C18 solid phase extraction cartridges (Agilent Bond Elut C18, 1 g bed mass, 6 mL) were equilibrated by flowing through 12 mL of methanol, followed by 12 mL of Milli-Q water. 50 mL of supernatant from a centrifuged bacterial culture was loaded into the cartridge, washed with 12 mL of Milli-Q water, and residual water was removed by a 30-second air pull. Indole was eluted from the cartridge with 3 mL of methanol, collected in 500 *μ*L fractions. The majority of indole was present in the first fractions, resulting in around 50-100x concentration.

Indole in the eluted fractions was quantified using QuantiChrom™ Indole Assay Kit (BioAssay Systems). 100 *μ*L of each fraction was added to individual wells of a 96-well plate, alongside a standard curve of 0 to 100 *μ*M indole in methanol. 100 *μ*L of QuantiChrom™ Indole Assay reagent was added to each well, shaken briefly, and absorbance read at 565 nm using a SpectraMax iD3 microplate reader (Molecular Devices). Absorbance values at 565 nm were also measured prior to the addition of QuantiChrom™ Indole Assay reagent, to identify increases in absorbance unrelated to indole concentration, and appropriately subtracted from the final value. The concentration of indole in each fraction was inferred from the absorbance value relative to the absorbance values of the standard curve. The limit of detection of this protocol was approximately 0.1 *μ*M indole in the original bacterial supernatant.

### SERS substrate (MLagg) preparation

Monolayer aggregates were prepared for the SERS signal enhancement. 500 *μ*L each of chloroform (CHCl_3_) and commercial (BBI Solutions) citrate-capped 80 nm gold nanoparticles (AuNPs) were added to an Eppendorf tube. 100 *μ*L of 1 mM cucurbit[5]uril (CB[5]) solution was then added and shaken for 1 min to initiate aggregation. The mixture was left to settle for the immiscible CHCl_3_ and aqueous phases to separate and the aggregated AuNPs to move to the phase interfaces (chloroform-aqueous and aqueous-air). The aqueous phase was washed with three 300 *μ*L aliquots of DI water to dilute the citrate salts and other supernatants, then concentrated by careful removal of the aqueous phase to form a 5 *μ*L aggregate droplet floating on the CHCl_3_. The droplet was deposited onto a pre-cleaned borosilicate glass coverslip (0.09 mm thick). Once dried, the resulting AuNP multilayer aggregate (MLagg) was rinsed with DI water and dried with N_2_. The MLaggs were oxygen plasma cleaned for 45 minutes (oxygen mass flow of 30 sccm, 90% RF power) using a plasma etcher (Diener electronic GmbH & Co. KG) to remove CB[5], citrate and other supernatants from the AuNP surfaces (verified using SERS). To re-introduce a scaffolding ligand, the MLaggs were immersed in 1 mM CB[5] solution prepared in 0.5 M HCl for 10 minutes, then rinsed with DI water and dried with N_2_. All chemicals were purchased from Sigma-Aldrich and aqueous solutions were prepared in deionized water (>18.2 MΩ cm^−1^, Purelab Ultra Scientific system).

### SERS measurements and pre-processing

SERS measurements were collected on a Renishaw InViva Raman confocal microscope, using a 20x objective (NA = 0.4) and 1200 lines mm^−1^ grating. A 785 nm excitation laser was used in extended scan mode, with 1 s integration time and 0.5% laser power (2.2 mW at the sample). All measurements were taken at room temperature, unless otherwise specified, and the spectra were calibrated with respect to Si. SERS measurements were taken for each MLagg substrate used for all assays, prior to immersing the MLagg in the supernatant solution. 400 *μ*L of the supernatant solution was pipetted into a polystyrene 96 well plate and the MLagg placed face down in the solution. It was ensured the MLagg was in contact with the solution in the well. The sample was left for 1 hour for molecules to diffuse to the hotspots and the sample to dry, then SERS measurements were taken at 10 spots on the MLagg and averaged (ten technical replicates).

All SERS spectra presented here were pre-processed using background correction and substrate normalisation, to ensure comparability across assays. Background correction was performed by iteratively fitting a polynomial to the base of the peaks. Substrate normalisation was performed relative to the cucurbit[5]uril (CB[5]) scaffold used in each of the SERS measurements, to account for potential variations in laser intensity and substrate enhancement factors at the time of measurement. All SERS spectra were normalised to the characteristic CB[5] peak at 829 cm^−1^, corresponding to the breathing mode of the macrocycle. This peak was assigned a value of 1, and all other intensities were scaled relative to this peak. Spectra presented in this form are referred to as ‘CB-normalised’ in Results. In addition, the reference CB[5] spectrum, acquired immediately prior to each SERS assay, was subtracted from the sample spectrum(‘CB-subtracted’). This subtraction was performed to show the contributions of the analytes from those intrinsic to the CB[5] scaffold, enabling direct comparison of spectral features arising from the biological samples alone.

### Semipolar metabolite analysis using LC-MS

Semi-polar metabolite profiling was performed by MS-Omics (Vedbak, Denmark). The analysis was carried out using a Vanquish LC (Thermo Fisher Scientific) coupled to a Orbitrap Exploris 240 MS (Thermo Fisher Scientific). The UHPLC used an adapted method described by Doneanu et al. (UPLC/MS Monitoring of Water-Soluble Vitamin Bs in Cell Culture Media in Minutes, Water Application note 2011, 720004042en). An electrospray ionization interface was used as ionization source. Analysis was performed in positive and negative ionization mode under polarity switching.

Untargeted data processing Metabolomics processing was performed untargeted using Compound Discoverer 3.3 (Thermo Fisher Scientific) and Skyline (24.1, MacCoss Lab Software) for peak picking and feature grouping, followed by a in-house annotation and curation pipeline written in MatLab (2022b, MathWorks). Identification of compounds were performed at four levels; Level 1: identification by retention times (compared against in-house authentic standards), accurate mass (with an accepted deviation of 3ppm), and MS/MS spectra, Level 2a: identification by retention times (compared against in-house authentic standards), accurate mass (with an accepted deviation of 3ppm). Level 2b: identification by accurate mass (with an accepted deviation of 3ppm), and MS/MS spectra, Level 3: identification by accurate mass alone (with an accepted deviation of 3ppm). Annotations on level 2b are based on accurate mass and MS/MS spectra measured with high resolution Orbitrap ESI-MS in mzCloud (Thermo Fisher Scientific), MassBank of North America (UC Davis) and the European MassBank (Helmholtz Centre for Environmental Research Leipzig). The annotations on level 3 are based on searches in the following libraries: *E. coli* metabolome database.

Metabolomic profiling yielded a total of 914 metabolites (144 annotated / known, 770 unannotated / unknown) across all assays. To quantify differences in metabolite abundance attributable to TnaA, the log_2_ fold-change (ΔMS) between WT and KO assays was computed. For each metabolite, the normalised peak area intensity in WT assays was divided by that in the corresponding KO assay, and the resulting ratio expressed as log_2_(WT/KO). Positive ΔMS values indicate enrichment in WT samples (i.e. metabolites associated with TnaA activity), whereas negative values indicate depletion relative to KO. Only metabolites exceeding the detection threshold (>3*σ* above background) were included in the analysis. This approach enabled systematic comparison of the metabolic profiles of WT and KO strains and identification of TnaA-dependent metabolites.

## Analytical techniques

The PCA uses an initial vector of size (1200, 4060) utilising three types of assays (enzymatic, cellular *E. coli* BW25113 and cellular *E. coli* 536), two types of strains (WT, or with TnaA, and KO, or without TnaA), 20 amino acids, and ten repeats for each spectrum recorded. This is reduced to (60, 1281) for Fig. 3 and is based on deducting KO spectra from WT spectra for each amino acid and using the 550cm^−1^ to 1700cm^−1^ wavenumber region.

To facilitate a more relevant interpretation of the PCA, we rotated the coordinate system such that the ‘null’ region (defined as spectra with minimal variation between WT and KO samples) was centered at the origin and aligned with the horizontal axis. The centroid of this region was translated to the origin, and subsequently the datapoints rotated to align the principal axis of the null region with the x-axis. This transformation yielded rotated components (PC1*, PC2*), preserving the relative geometry of all points.

In order to try and fully identify metabolite signatures in the spectral data, we utilised various dimensionality reduction techniques. We find PCA alone provided a more relevant and interpretable representation than t-SNE or UMAP methods for SERS data (Fig. 2), particularly given the need to quantify the loadings (wavenumbers of interest) and PCA’s deterministic approach. We also utilised a two-step analytical procedure, performing low-rank PCA and then embedding these components with UMAP. This provided cleaner clusters than PCA alone, albeit with less intuitive interpretability. SI Fig. 3.7 shows results for the UMAP technique applied on WT minus KO SERS data, as shown in Fig. 2.

## Author contributions statement

M.V., A.Z., K.A., D.F. and J.J.B. designed the experiments. M.V. prepared all bacteria-related assays and provided the bacterial samples for SERS. A.Z. supplied both bacteria strains including the genetically modified strains. C.C. generated the 536 Δ*tnaA* strain. Enzymatic assays were prepared by K.A. E.W.W. fabricated MLagg substrates and carried out SERS measurements, with advice from M.K. and T.F.K. Data analysis was performed by M.V., E.W.W, D.F. and J.J.B. Manuscript drafting was led by M.V. and all authors contributed to data interpretation, figures and manuscript editing.

## Acknowledgments

The authors would like to thank Dr. Isabel Askenasy for their assistance with protein purification and for their helpful comments and discussions. A.Z. is a recipient of a Transition To Independence (TTI) fellowship from the School of Biological Sciences at the University of Cambridge and was supported by funding from the Rosetrees Trust (JS16/TTI2021/1), the Isaac Newton Trust (21.22(a)iii) and the School of Biological Sciences at the University of Cambridge. J.J.B is funded by the European Research Council (ERC) under Horizon 2020 research and innovation programme PICOFORCE (Grant Agreement No. 883703), POSEIDON (Grant Agreement No. 861950) and EPSRC (Cambridge NanoDTC EP/L015978/1, EP/L027151/1, EP/X037770/1).

## Data availability

The data that support the findings of this study are available upon request.

