## Supplementary Information for "Nanoplasmonic SERS-enabled structural identification of an uncharacterised indole metabolite directly in E. coli cultures"

### Table of Contents

**Supplementary Fig. 1.1: Spiking of cultures with indole** 3

**Supplementary Fig. 1.2: SERS of cellular assays without amino acid supplementation** 4

**Supplementary Fig. 1.3: DFT-calculated spectra of indole and its mono-oxidised and protonated forms** 5

**Supplementary Fig. 1.4: SERS measurements of indole at 100nM concentration** 6

**Supplementary Fig. 1.5: Effect of different culture media on SERS spectra of *E. coli*** 7

**Supplementary Fig. 1.6: SERS of purified TnaA enzyme only** 8

**Supplementary Fig. 2.1: Profiling of *E. coli* BW25113 cellular assays with adjustment by assay without amino acid supplementation** 9

**Supplementary Fig. 2.2: Profiling of *E. coli* 536 cellular assays with adjustment by assay without amino acid supplementation** 10

**Supplementary Fig. 2.3: Higher resolution of WT minus KO spectra for cellular assays** 11

**Supplementary Fig. 2.3 (cont'd): Higher resolution of WT minus KO spectra for cellular assays** 12

**Supplementary Fig. 2.4: Profiling of *E. coli* BW25113 cellular assays with adjustment by assay without cells and amino acid only.** 13

**Supplementary Fig. 2.5: Profiling of *E. coli* BW25113 cellular assays with adjustment by assay without cells and amino acid only.** 14

**Supplementary Fig. 2.6: All enzymatic and cellular *E. coli* BW25113 assays.** 15

**Supplementary Fig. 2.7: Repeat assays for *E. coli* BW25113 WT and KO strains** 16

**Supplementary Fig. 2.8: PBS-washed assays versus unwashed assays** 17

**Supplementary Fig. 3.1: PCA of WT minus KO spectra showing PC1-PC2, PC1-PC3 and PC2-PC3** 18

**Supplementary Fig. 3.2: PCA of WT, KO spectra by assay type** **19**

**Supplementary Fig. 3.3: PCA of WT, KO spectra by amino acid** **20**

**Supplementary Fig. 3.4: PCA of WT, KO, and amino acid-only spectra with points connected to indicate relative similarity** **21**

**Supplementary Fig. 3.5: PCA loadings of BW25113 supernatant WT minus KO spectra** **22**

**Supplementary Fig. 3.6: Rotated PCA for amino acid-supplemented cultures** **23**

**Supplementary Fig. 3.7: UMAP of WT minus KO spectra** **24**

**Supplementary Fig. 4.1: Analysis of LC-MS detected metabolites by $\log_2$ fold-change.** **26**

**Supplementary Fig. 5.1: Structural Equation Modelling Analysis of I*** **27**

**Supplementary Table 1: Composition of the base of artificial urine media** **29**

**Supplementary Table 2: Composition of the supplement solutions of artificial urine media** **29**

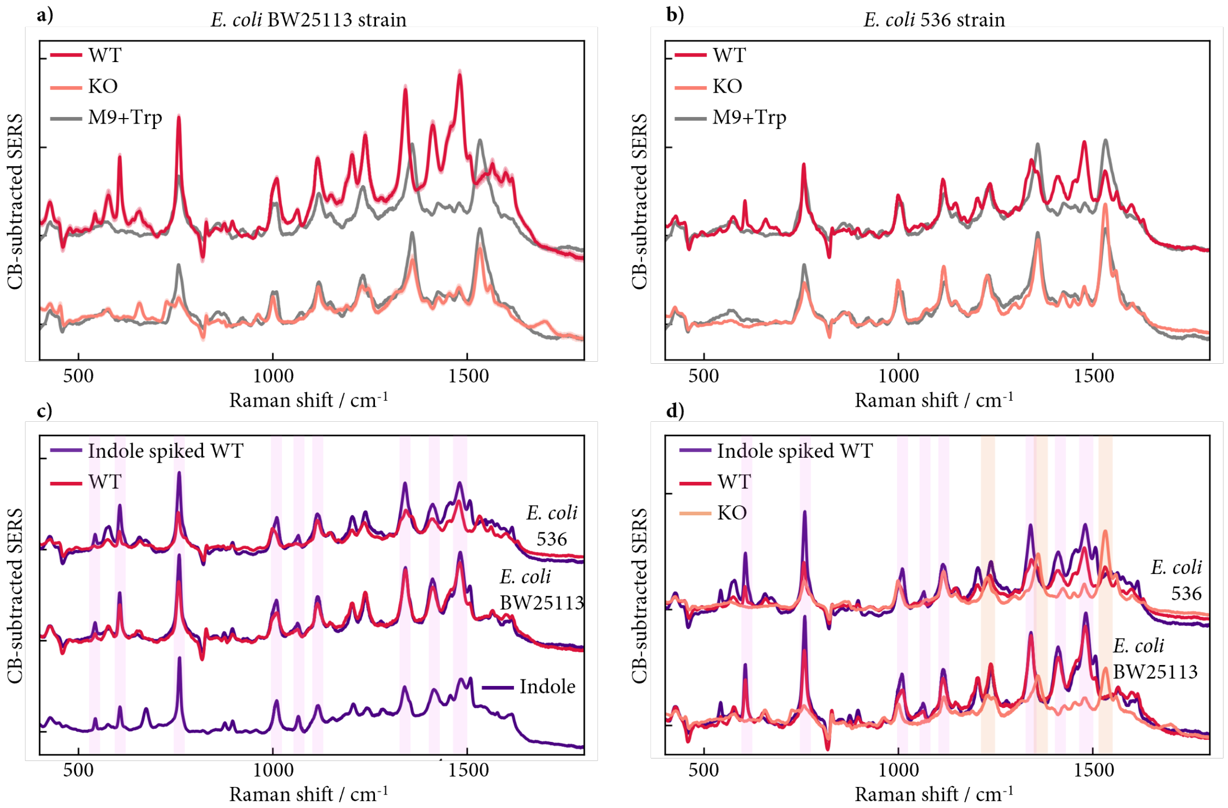


**Supplementary Fig. 1.1: Spiking of WT supernatant with pure indole (10mM)**. CB-subtracted SERS of WT, KO and culture media-only assays using M9 supplemented with Trp shown for (a) *E. coli* BW25113 and (b) *E. coli* 536 strains. (c) Comparison of subtracted SERS between unspiked WT (red) and indole-spiked WT (indigo) supernatant shows enhancement of peaks corresponding to pure indole SERS (lower spectrum). (d) SERS of unspiked WT, indole-spiked WT and unspiked KO strains showing indole peaks (pink shading) and undegraded Trp (orange shading) in KO strain.

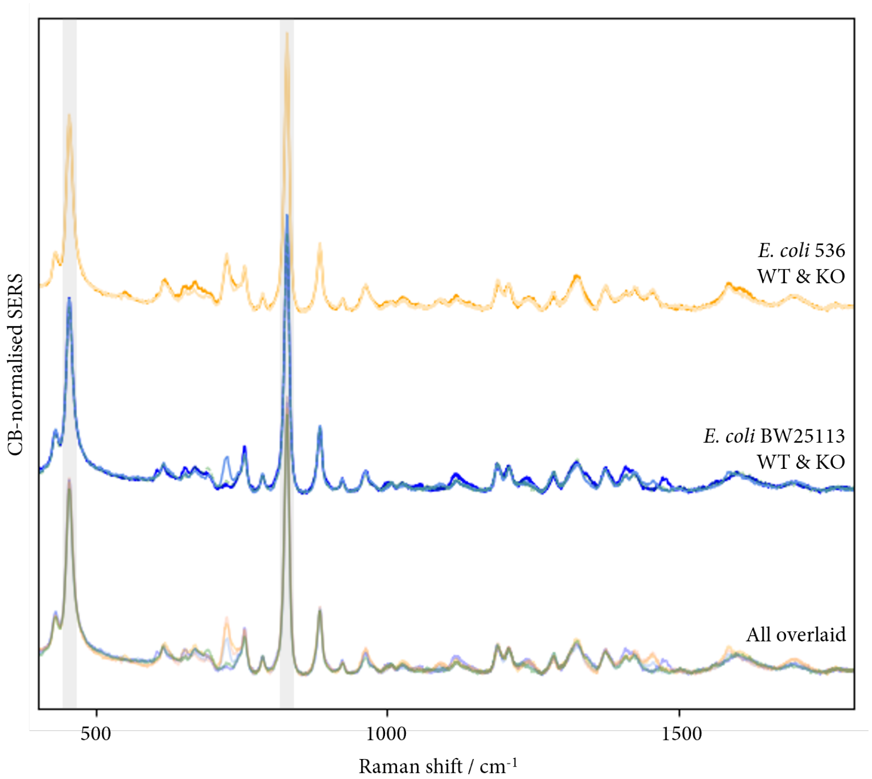


**Supplementary Fig. 1.2: SERS of cellular assays without amino acid supplementation.** CB-normalised SERS of WT and KO strains cultured in M9 salts only, without amino acid supplementation. Strong alignment between WT and KO for both *E. coli* 536 and *E. coli* BW25113 strains. KO strain for both *E. coli* 536 and *E. coli* BW25113 expressing a kanamycin resistance gene used as a marker to confirm the gene knockout method. Kanamycin resistance protein is intracellular, thus should not be present in the supernatant, as confirmed by the high degree of alignment between WT and KO strains in the absence of external amino acid supplementation. Grey regions highlight the peaks associated with CB[5], the scaffold in the SERS MLagg substrate used for all experiments, and common to all SERS spectra.

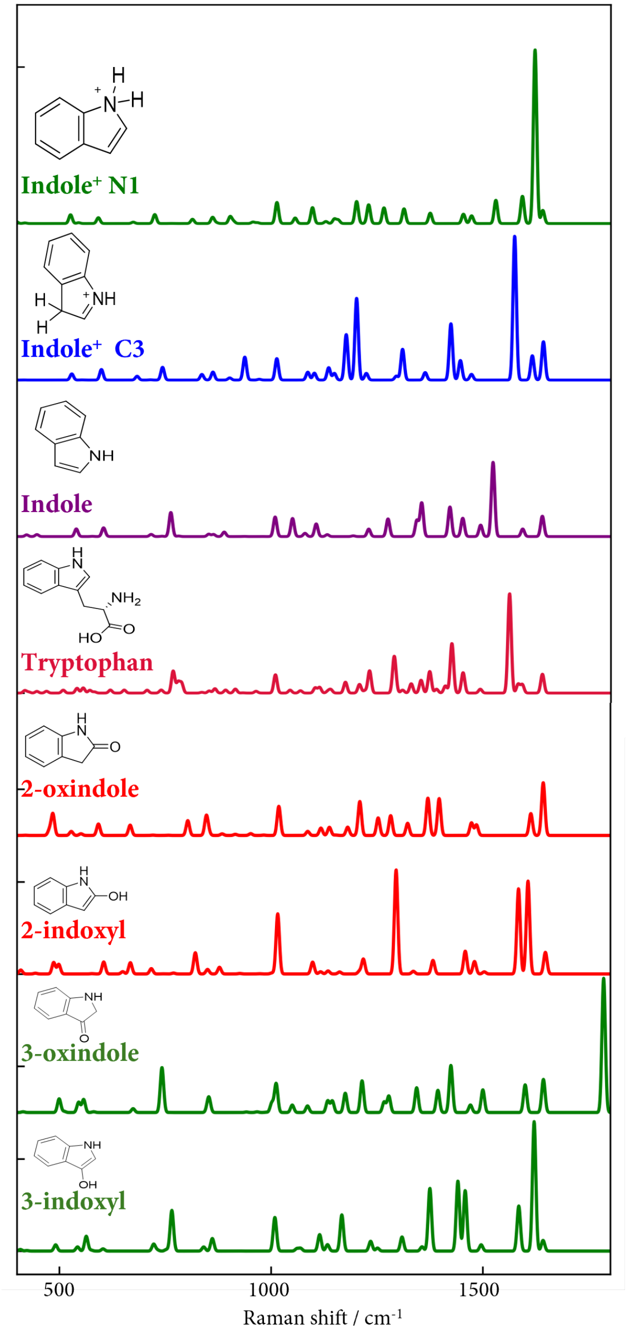


**Supplementary Fig. 1.3: Density functional theory (DFT)-derived spectra of indole, tryptophan and indole derivatives.** Protonated and mono-oxidised forms of indole shown for reference, neither of which precisely match I*. Scaling factor of 0.975 used.

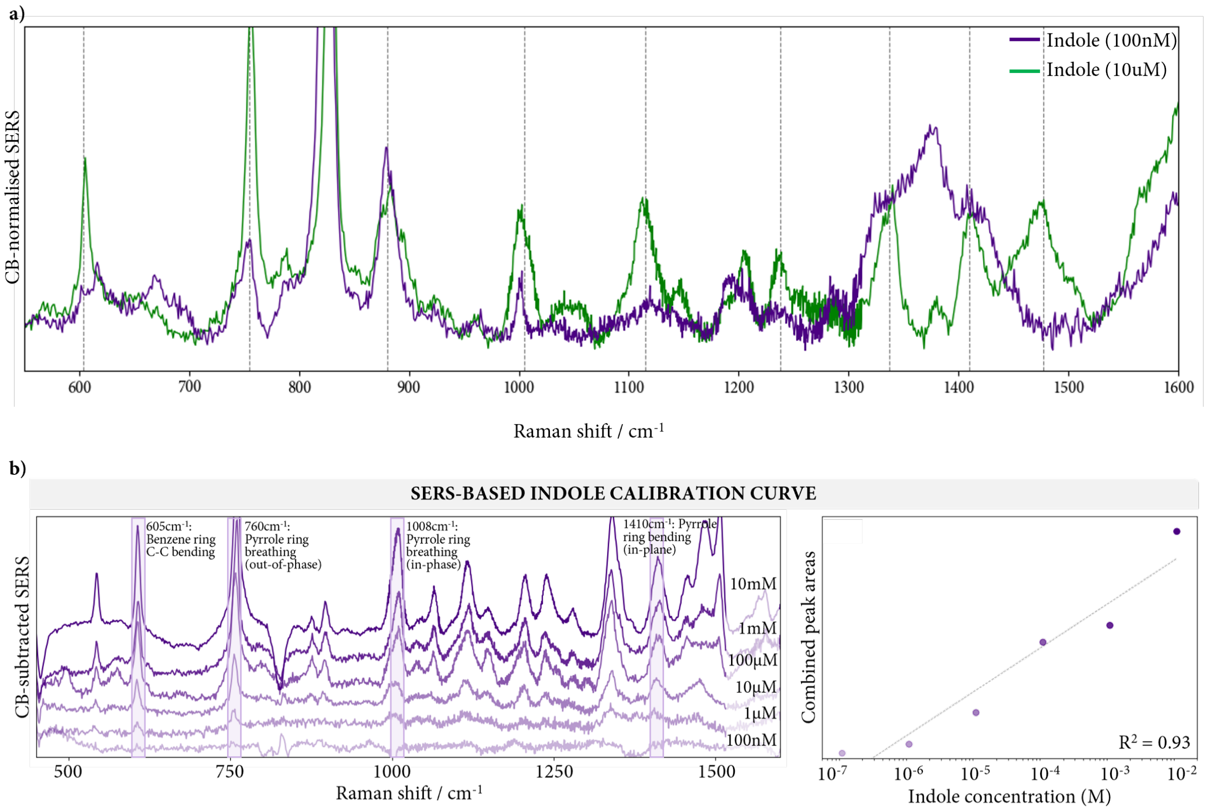


**Supplementary Fig. 1.4: SERS measurements of indole at 100nM concentration.** SERS of 10um (100x) rescaled and shown for peak references. Gray vertical lines indicate the indole band assignments as shown in Fig.1. Significant peak alignment observed between both spectra, except for the 1300-1500cm-1 range, where peak broadening is observed in the 100nM spectrum.

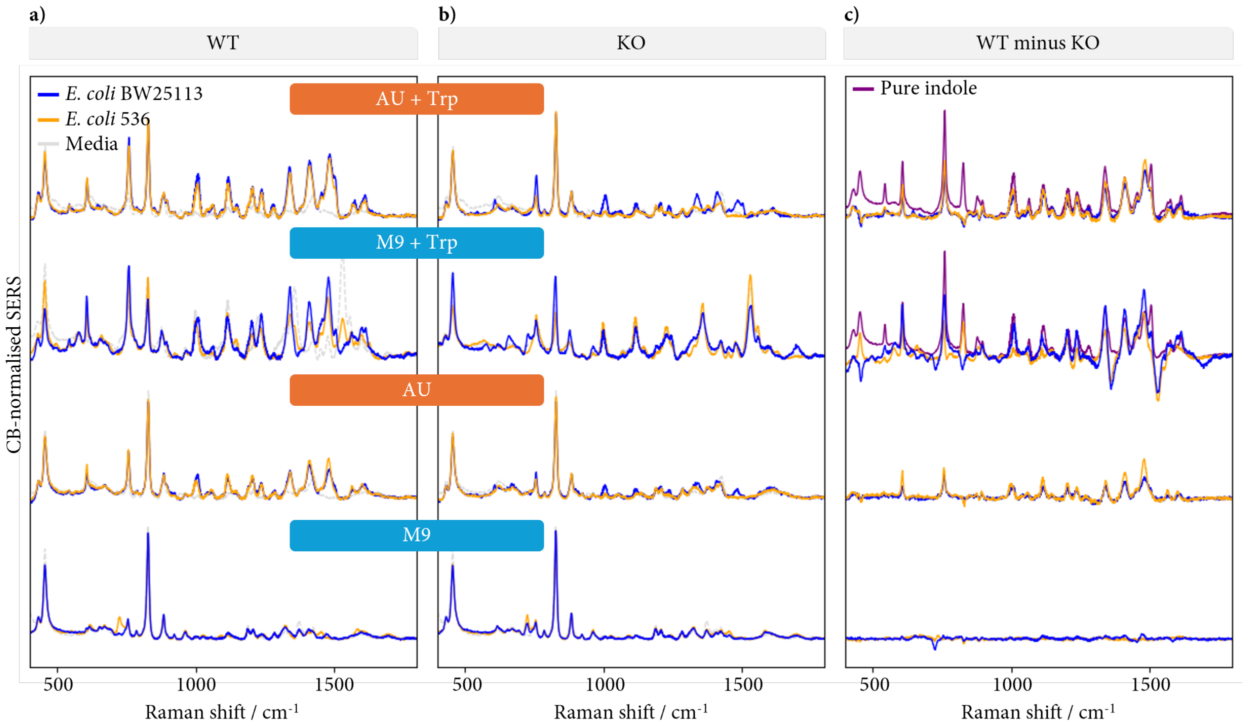


**Supplementary Fig. 1.5: Effect of different culture media on SERS spectra of *E. coli.*** Indole spectra isolated in more complex artificial urine media (AU), mimicking clinically relevant settings. (a, b) SERS for *E. coli* BW25113 (blue) and *E. coli* 536 (orange) (a) WT and (b) KO strains in AU and M9 with and without Trp supplementation. (c) WT spectrum minus KO spectrum showing well-aligned peaks in both media, with indole spectra superimposed for reference. Weak but significant SERS activity seen for both WT minus KO spectra in AU alone, indicating the presence of various amino acids contained in the AU media (see Supplementary Methods).

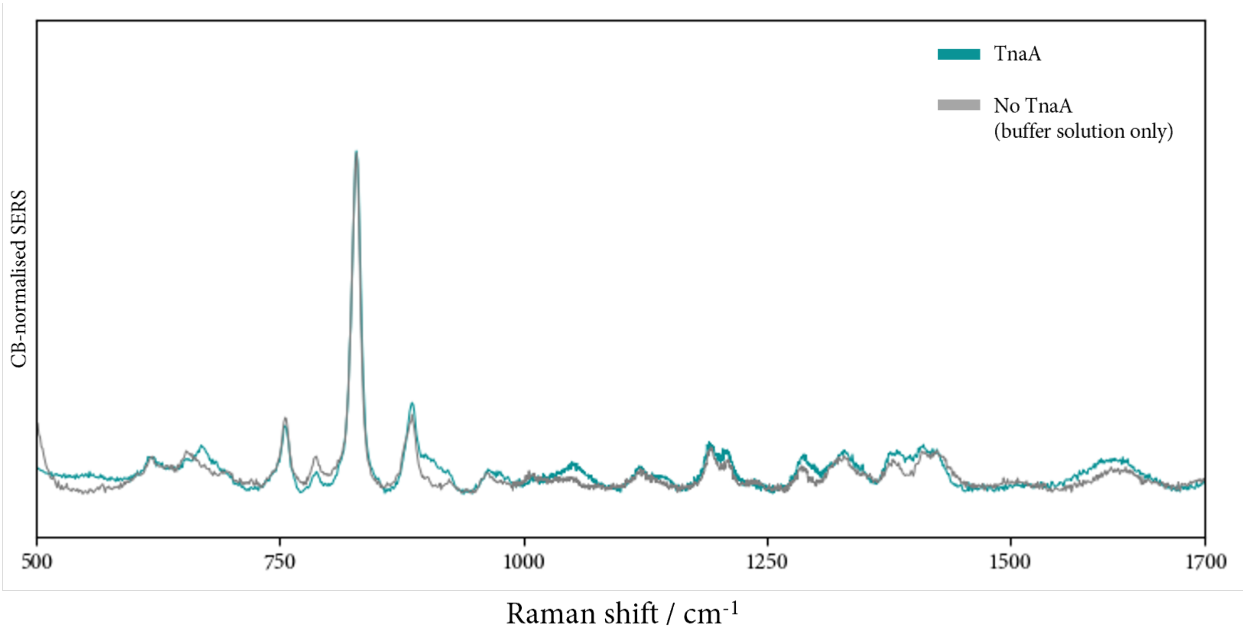


**Supplementary Fig. 1.6: SERS of purified TnaA enzyme only.** Comparison of SERS spectra of purified TnaA with TnaA-filtered buffer solution containing no TnaA enzyme. High degree of similarity in spectra, showing purified TnaA provided negligible SERS activity relative to the buffer solution. See SI Methods for further details.

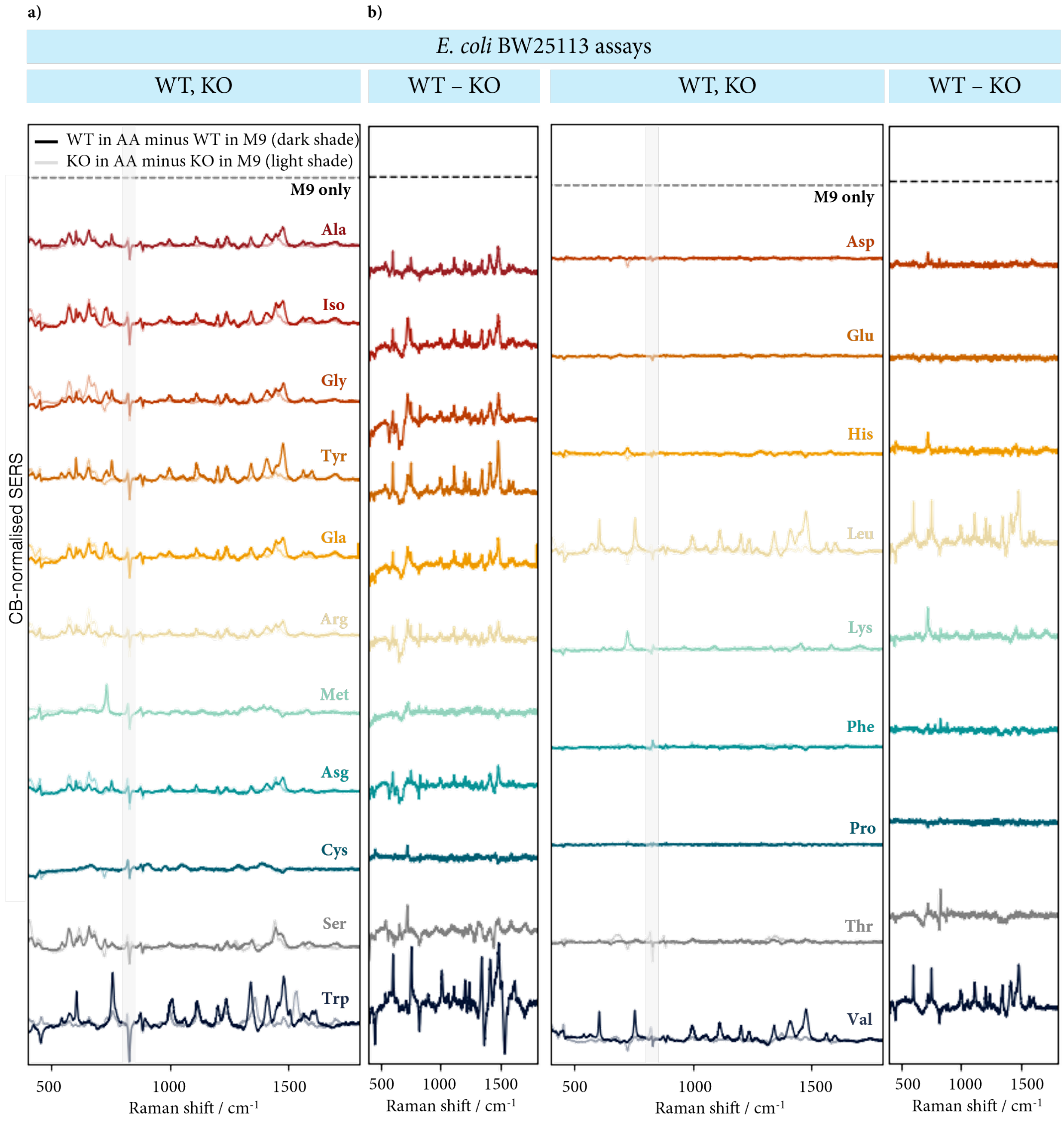


**Supplementary Fig. 2.1: Profiling of *E. coli* BW25113 cellular assays with adjustment by corresponding assay without amino acid supplementation and in M9 salts only.** (a,b) SERS of *E. coli* BW25113 for (a) WT and KO strains minus the assay in M9 only, and (b) WT – KO strains. Assays in M9 show minimal activity indicating the presence of the amino acid contributes to the different spectral profiles observed.

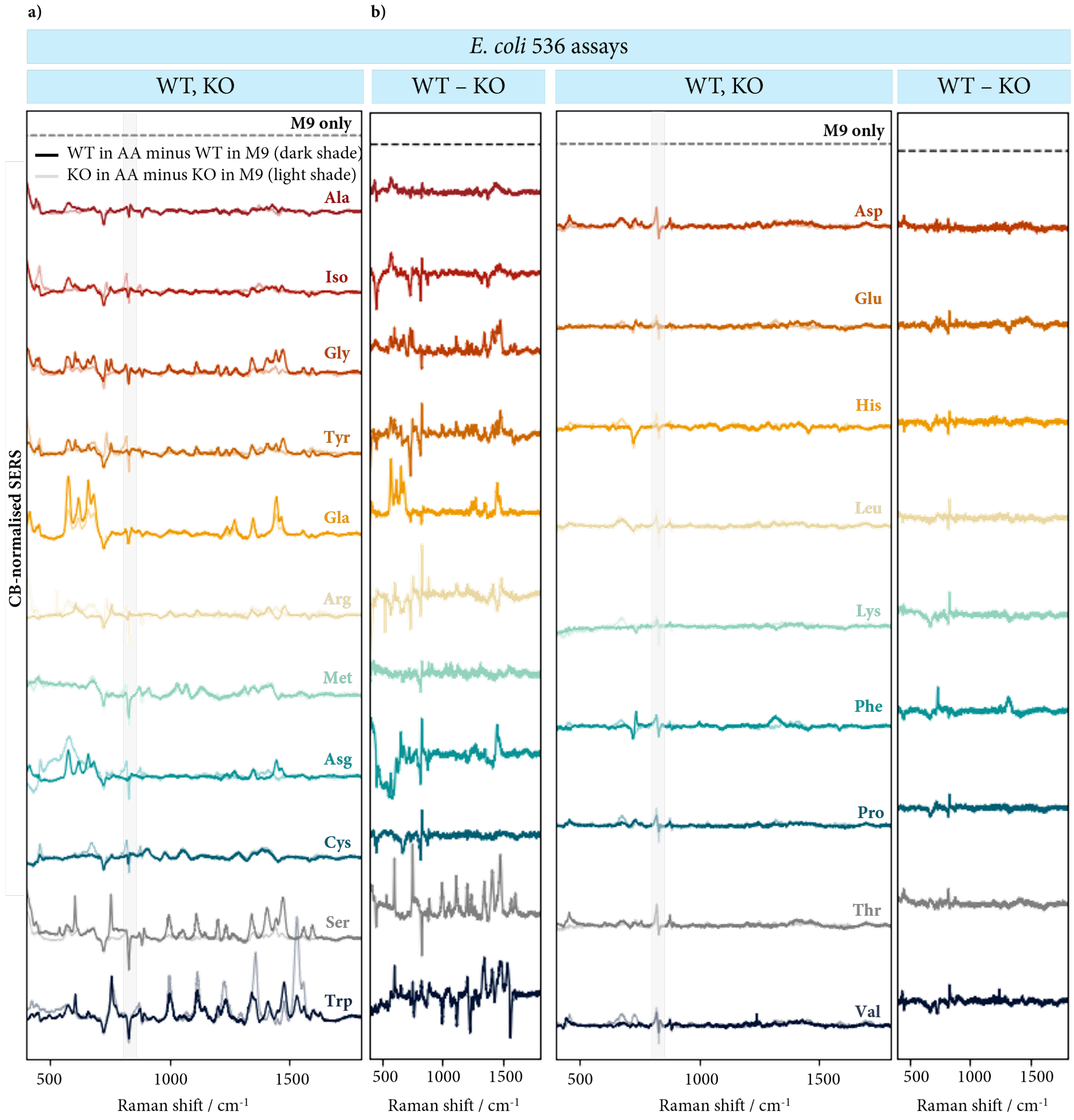


**Supplementary Fig. 2.2: Profiling of *E. coli* 536 cellular assays with adjustment by corresponding assay without amino acid supplementation and in M9 salts only.** (a,b) SERS of *E. coli* BW25113 for (a) WT and KO strains minus the assay in M9 only, and (b) WT – KO strains. Assays in M9 show minimal activity indicating the presence of the amino acid contributes to the different spectral profiles observed.

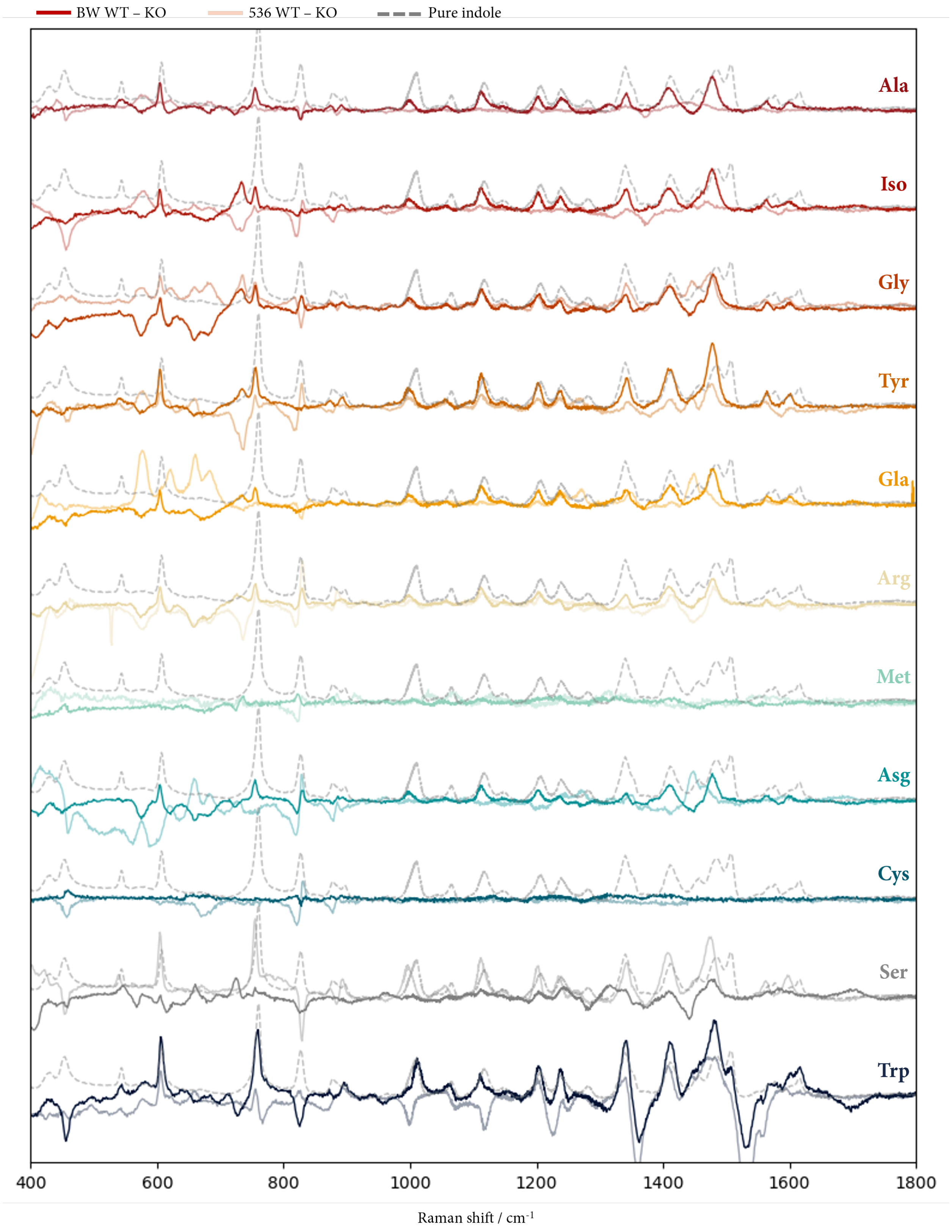


**Supplementary Fig. 2.3: Higher resolution of WT minus KO spectra for both *E. coli* BW25113 and *E. coli* 536 strains.** SERS for pure indole (10mM) superimposed to compare spectral differences.

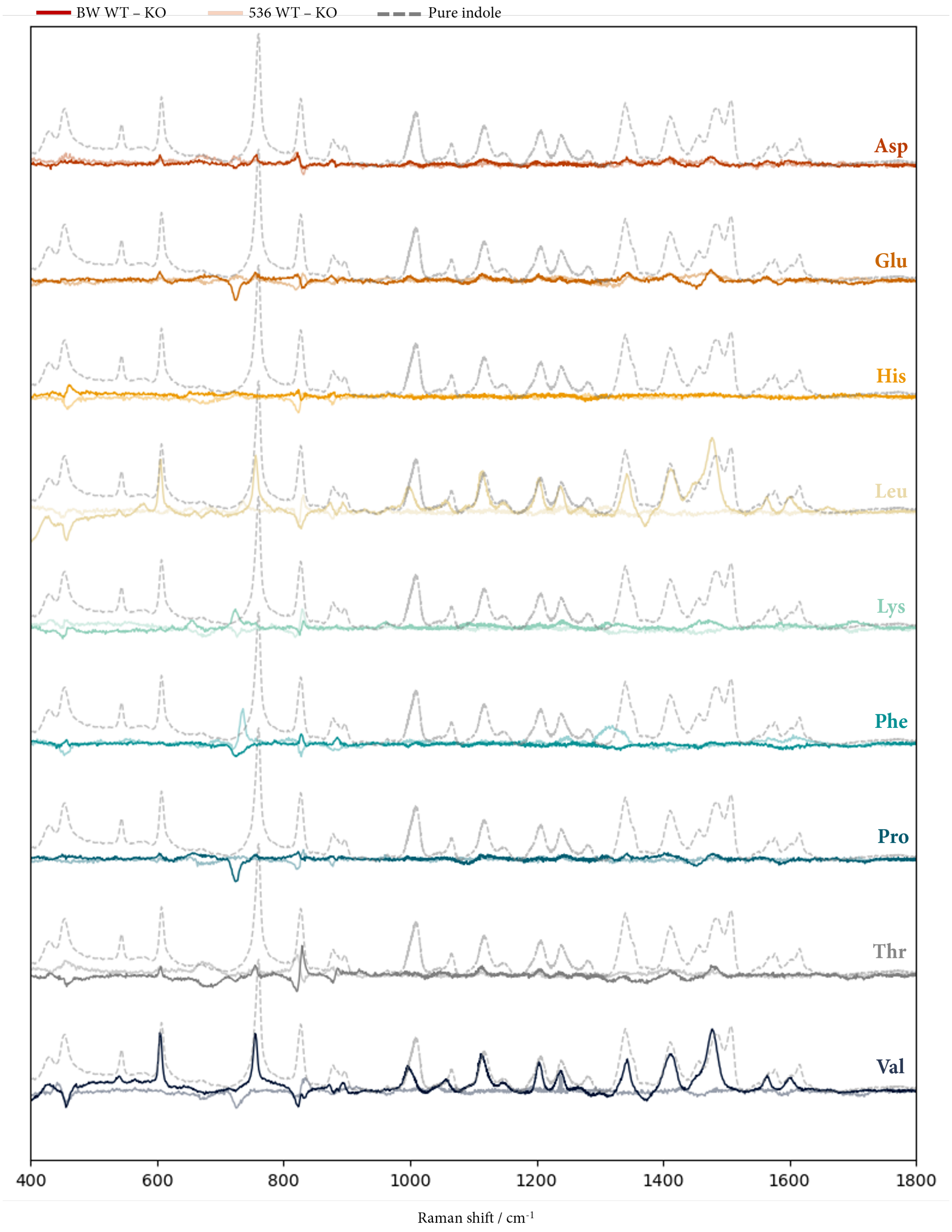


**Supplementary Fig. 2.3 (cont'd): Higher resolution of WT minus KO spectra for both *E. coli* BW25113 and *E. coli* 536 strains.** SERS for pure indole (10mM) superimposed to compare spectral differences.

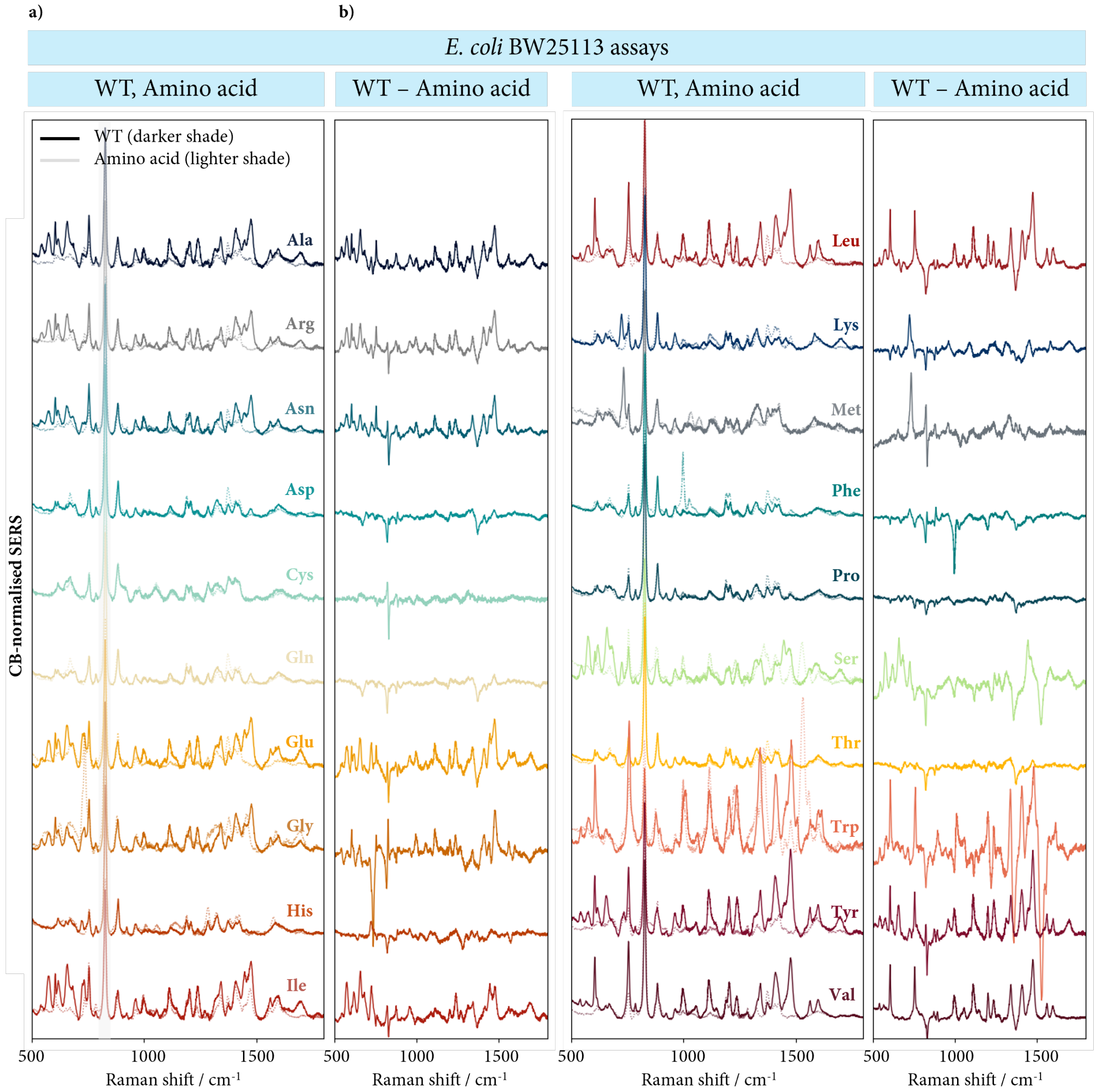


**Supplementary Fig. 2.4: Profiling of *E. coli* BW25113 cellular assays with adjustment by assay without cells and amino acid only.** (a,b) SERS of *E. coli* BW25113 for (a) WT strain and amino acid controls and (b) WT – amino acid control. Significant differences in spectra indicate the I* signature is not due to the amino acid controls and is a function of the cellular assay cultured in the amino acid.

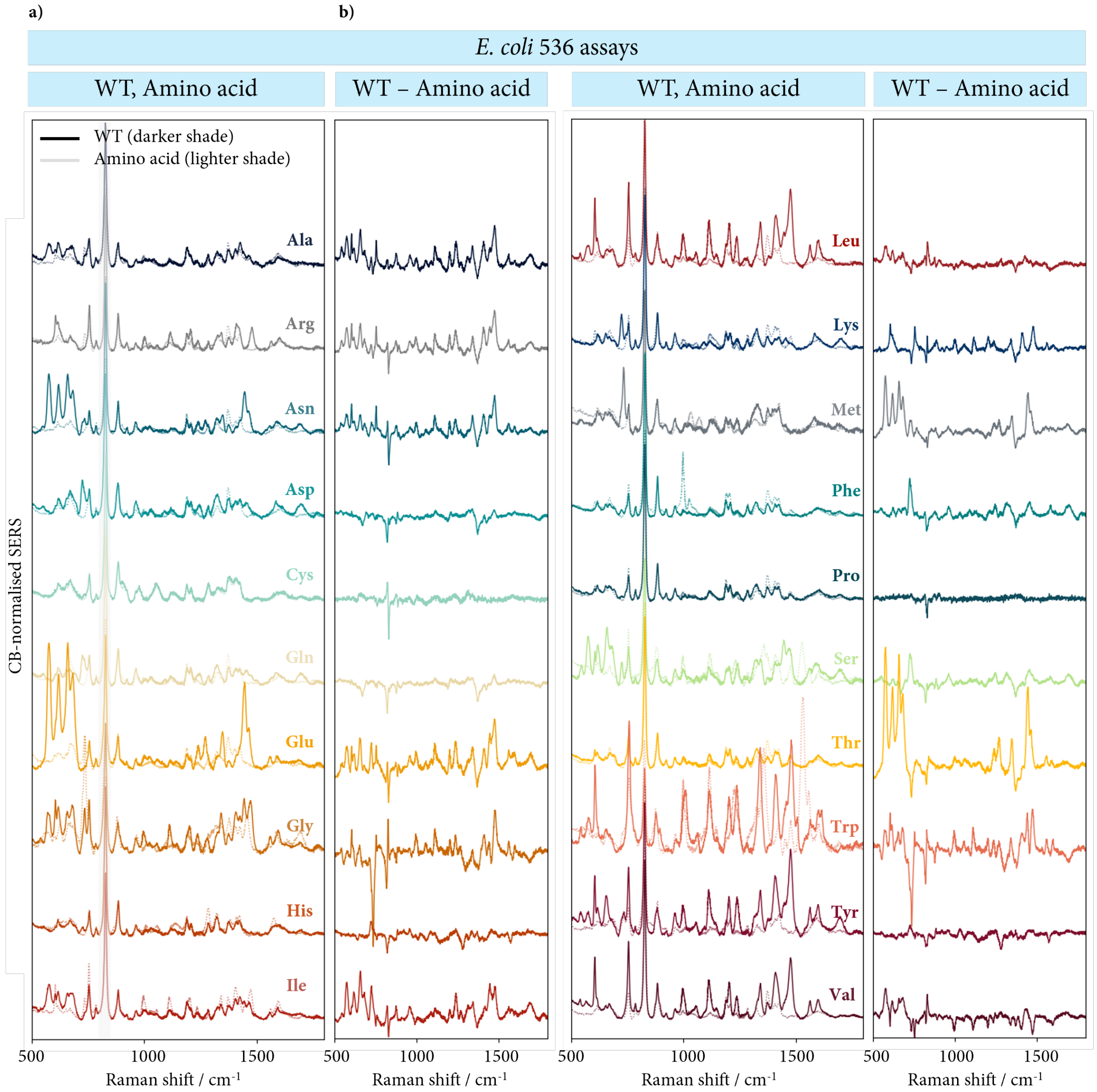


**Supplementary Fig. 2.5: Profiling of *E. coli* 536 cellular assays with adjustment by assay without cells and amino acid only.** (a,b) SERS of *E. coli* BW25113 for (a) WT strain and amino acid controls and (b) WT – amino acid control. Significant differences in spectra indicate the I* signature is not due to the amino acid controls and is a function of the cellular assay cultured in the amino acid.

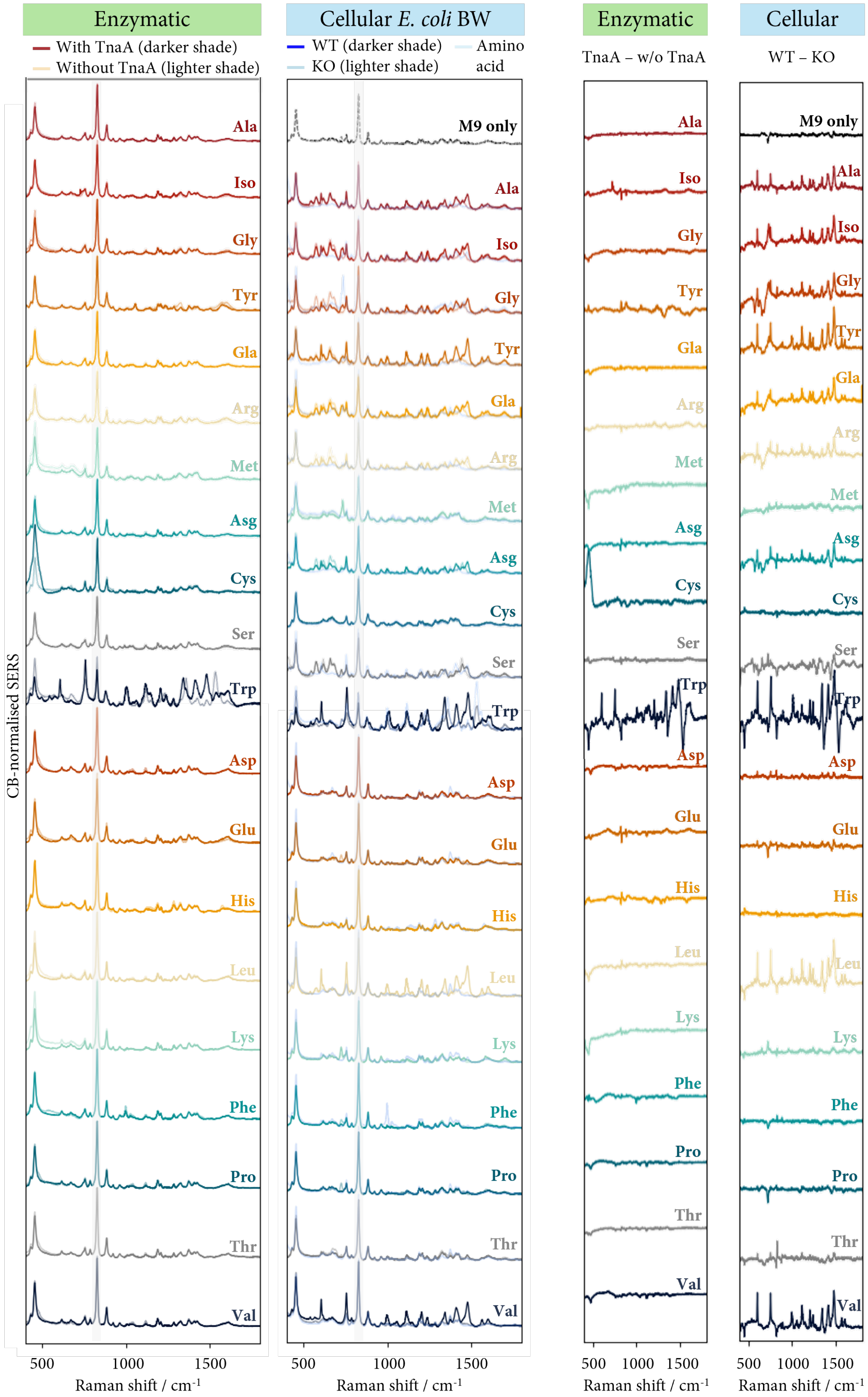


**Supplementary Fig. 2.6: All spectra shown for both enzymatic and cellular *E. coli* BW25113 assays.**

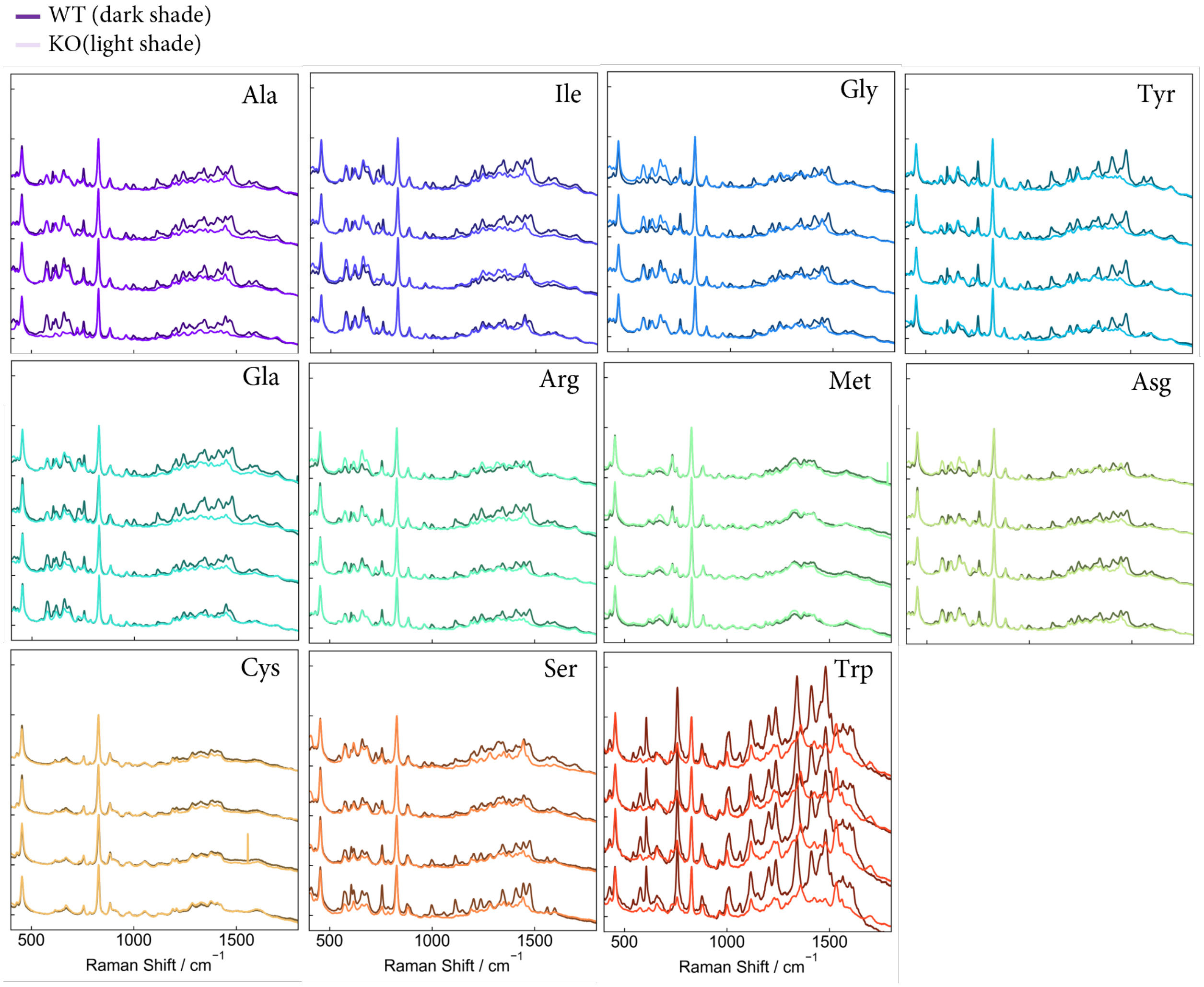


**Supplementary Fig. 2.7: SERS of biological repeat assays for *E. coli* BW25113 WT and KO strains.** Four biological repeats and ten technical repeats performed. SERS measurements averaged across ten technical repeats. Consistent spectral profile seen across all assays for the same amino acid.

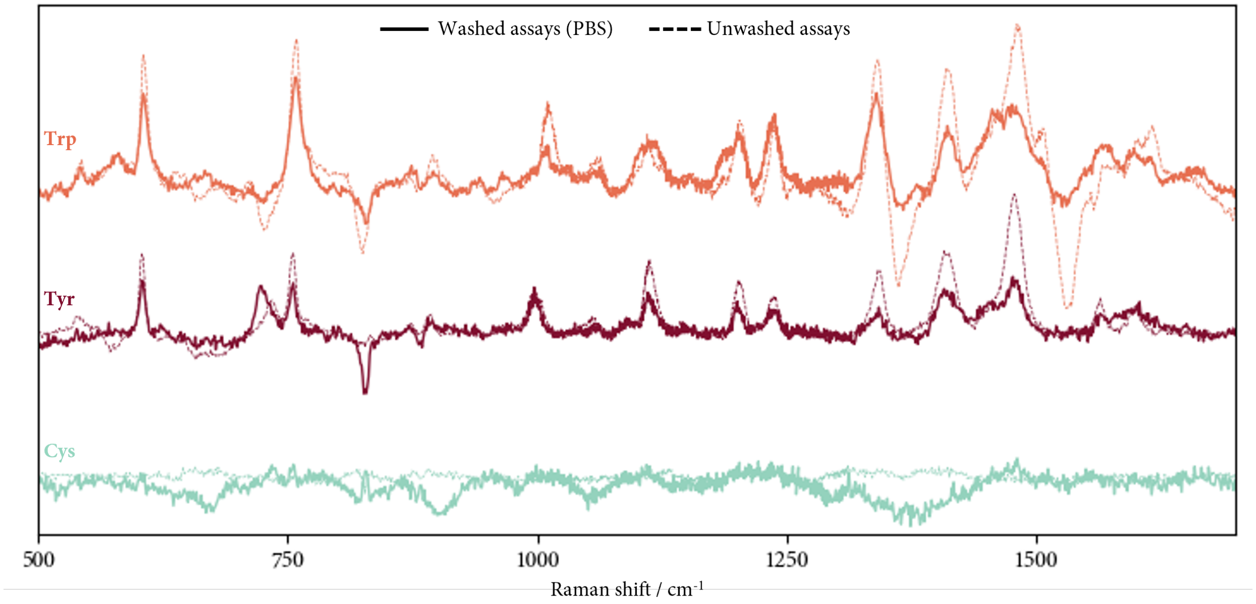


**Supplementary Fig. 2.8: PBS-washed versus unwashed WT minus KO spectra for selected *E. coli* BW25113 assays.** Tryptophan-supplemented, tyrosine-supplemented and cysteine-supplemented *E. coli* BW25113 supernatant assays shown as they represent indole, I*, and no I* signatures, respectively. Washed WT minus KO spectra resemble corresponding unwashed assay spectra, suggesting a minimal indole carryover effect. See Fig. 2 and Methods in the main text for more details.

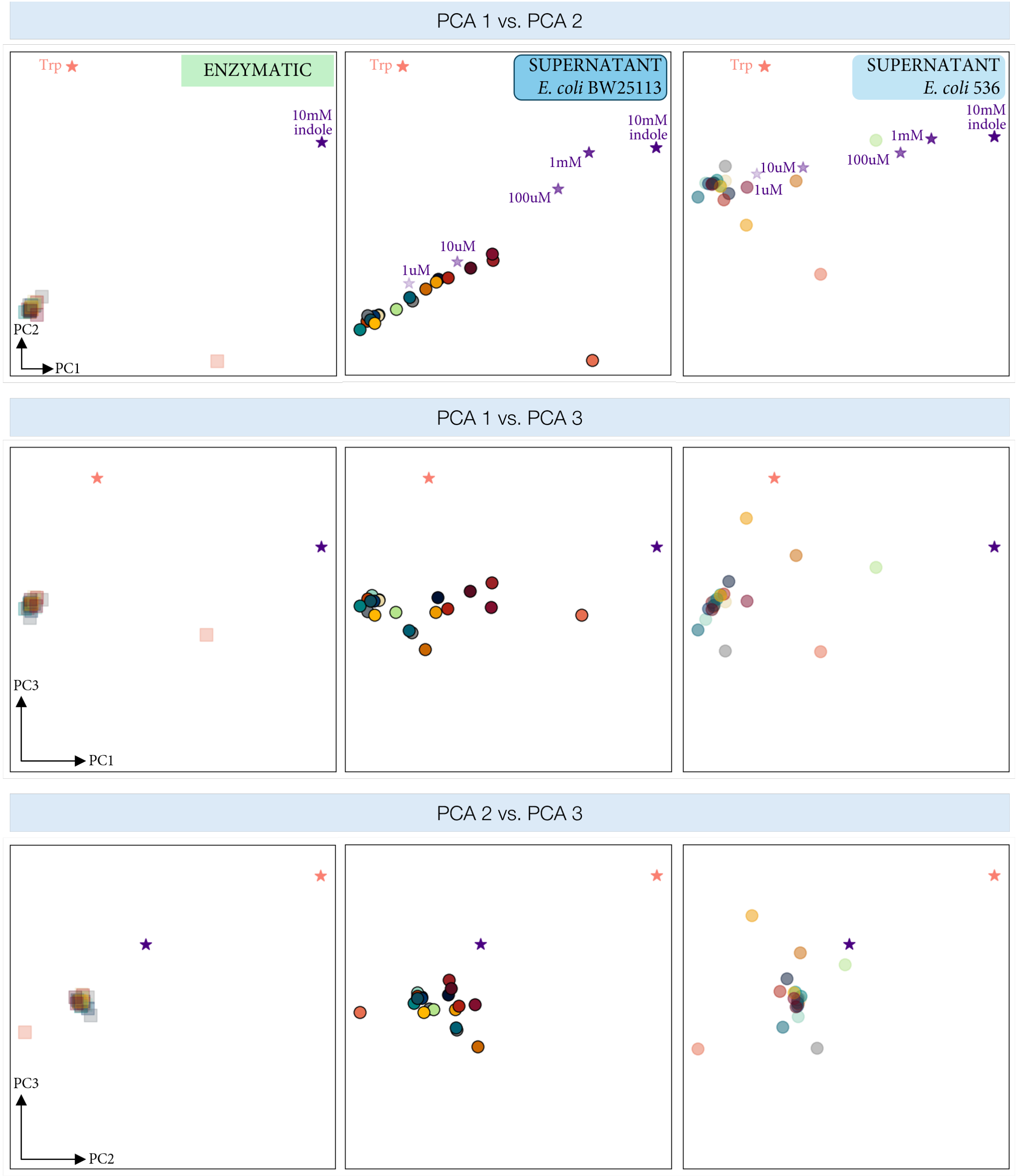


**Supplementary Fig. 3.1: PCA of WT minus KO spectra for enzymatic, *E. coli* BW25113 and *E. coli* 536 assays.** Plot of PC1-PC2 indicates WT minus KO assays lie on a line. PC1-PC3 and PC2-PC3 plots indicate this line may lie on a 2D plane, given the lack of meaningful subsequent components in discriminating the data. First two components explain c.70% of the variation.

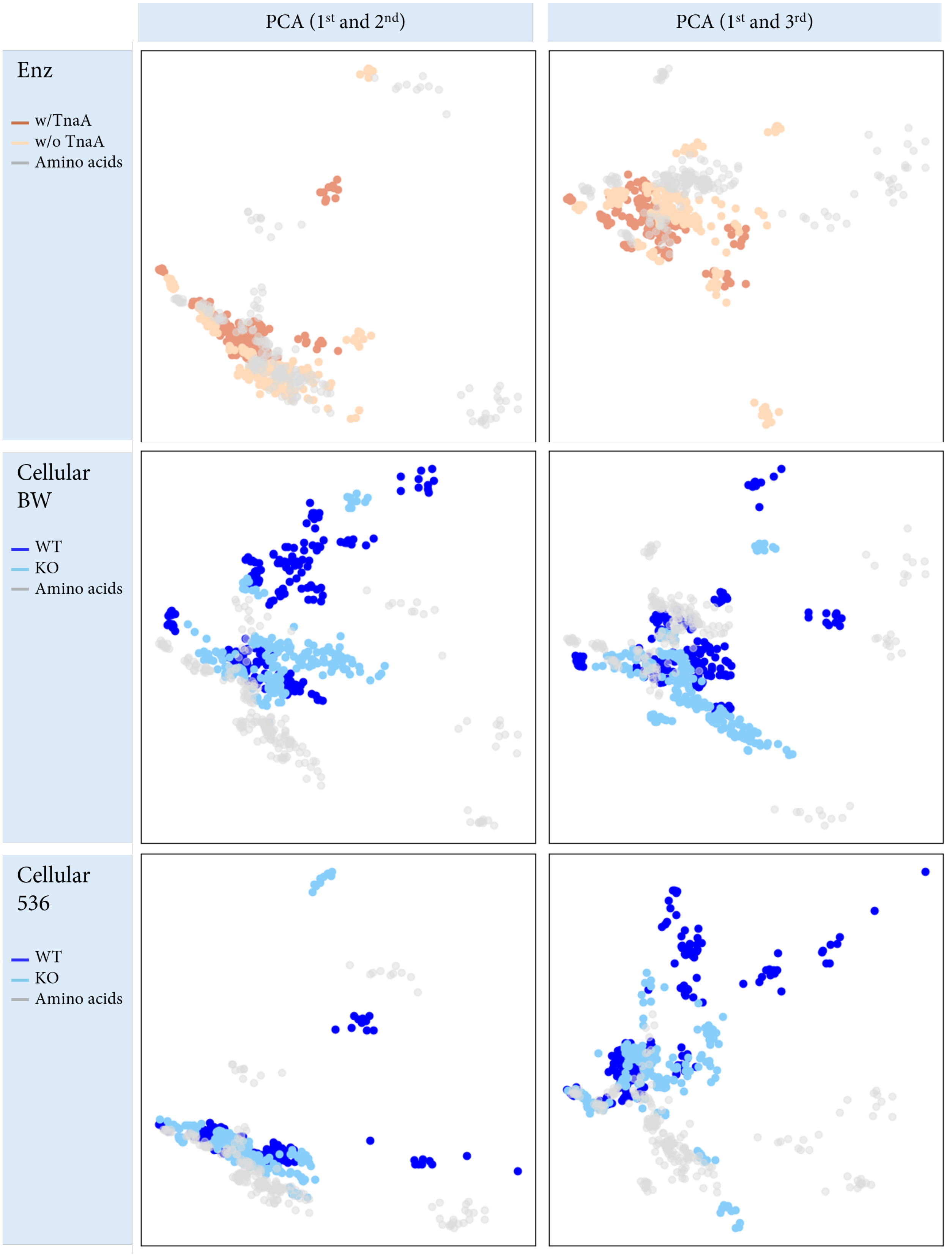


**Supplementary Fig. 3.2: PCA of WT and KO spectra for enzymatic, *E. coli* BW25113 and *E. coli* 536 assays, coloured by assay type.** PC1-PC2 and PC1-PC3 shown for reference.

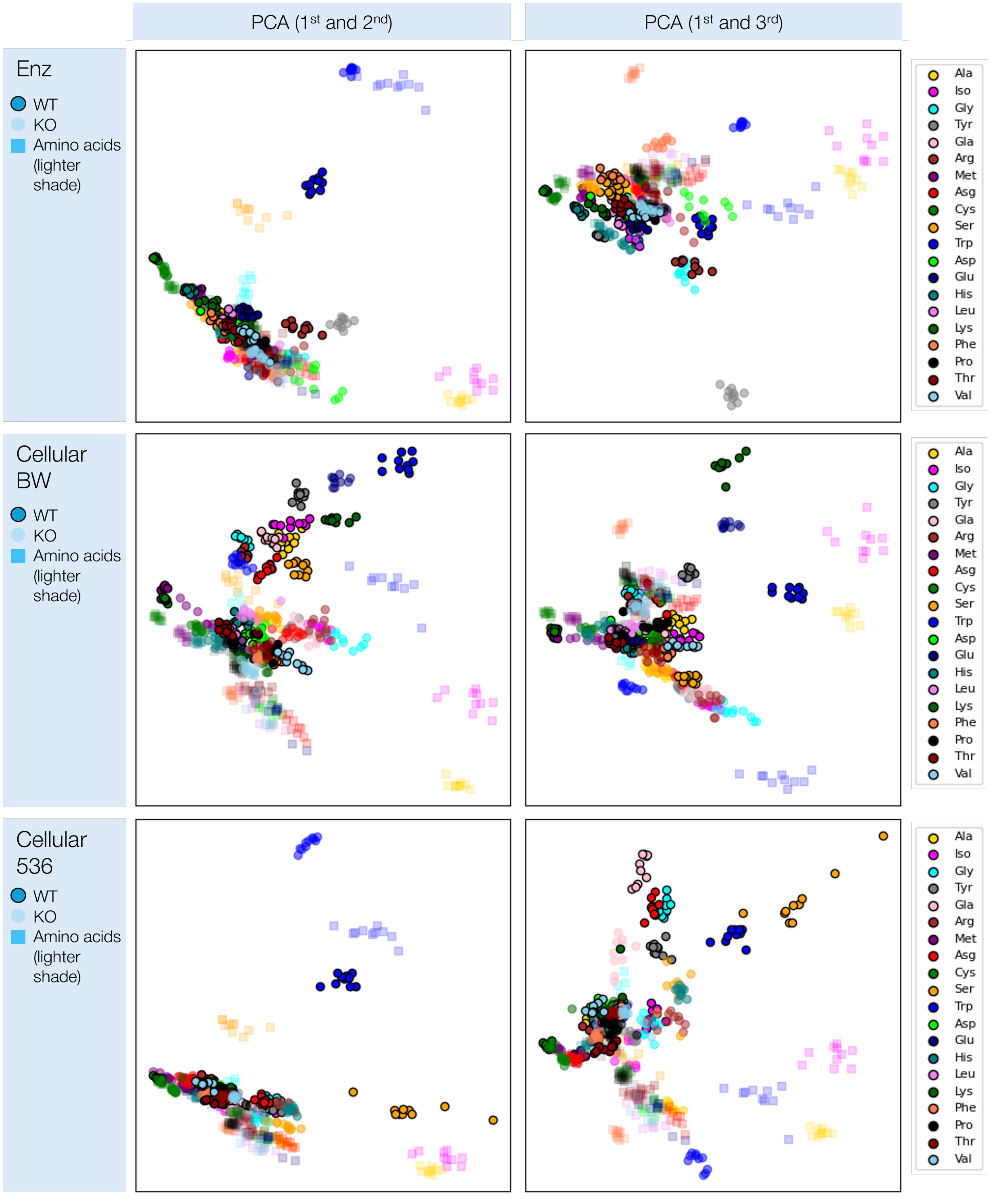


**Supplementary Fig. 3.3: PCA of WT and KO spectra for enzymatic, *E. coli* BW25113 and *E. coli* 536 assays, coloured by amino acid.** PC1-PC2 and PC1-PC3 shown for reference.

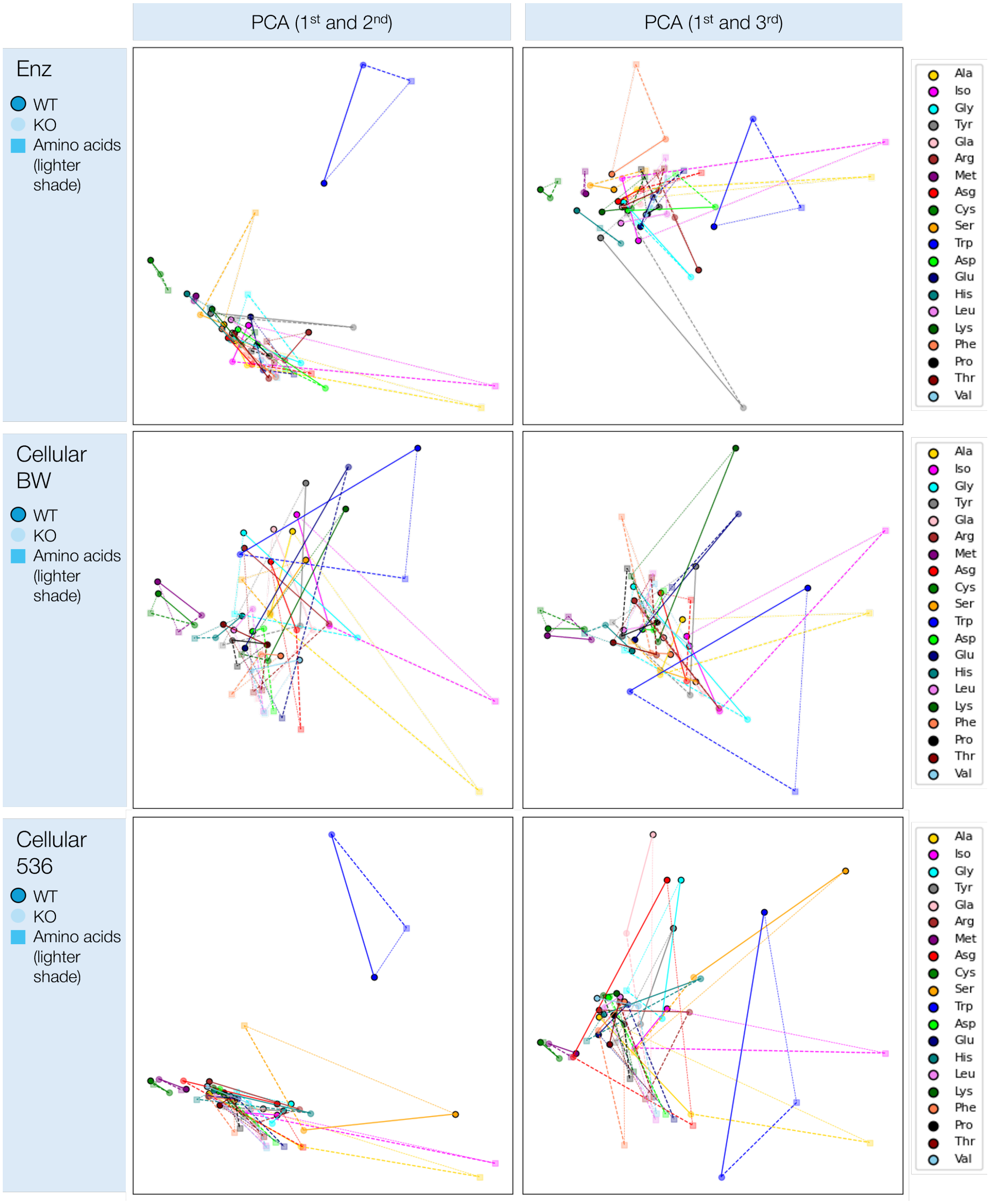


**Supplementary Fig. 3.4: PCA of WT, KO and amino acid only spectra for enzymatic, *E. coli* BW25113 and *E. coli* 536 assays, coloured by amino acid.** PC1-PC2 and PC1-PC3 shown for reference. Clear difference seen in amino acid only spectra and WT and KO spectra as shown by the relative distance between points.

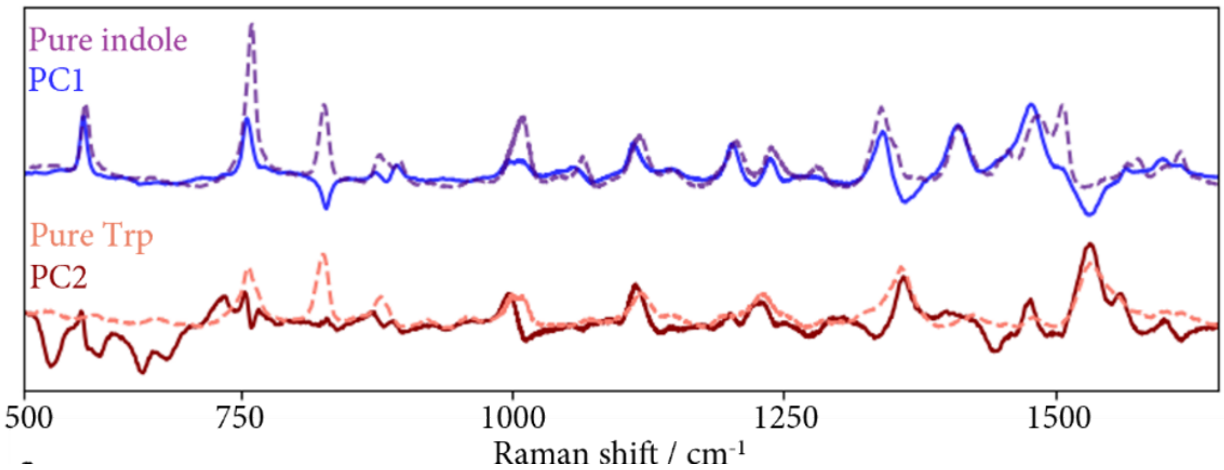


**Supplementary Fig. 3.5: PCA loadings from PCA of supernatant WT minus KO spectra for *E. coli* BW25113 strain, as shown in Fig. 3b).** PC1 and PC2 weightings (solid), compared to free indole and pure tryptophan spectra (dashed), showing a high degree of spectral alignment.

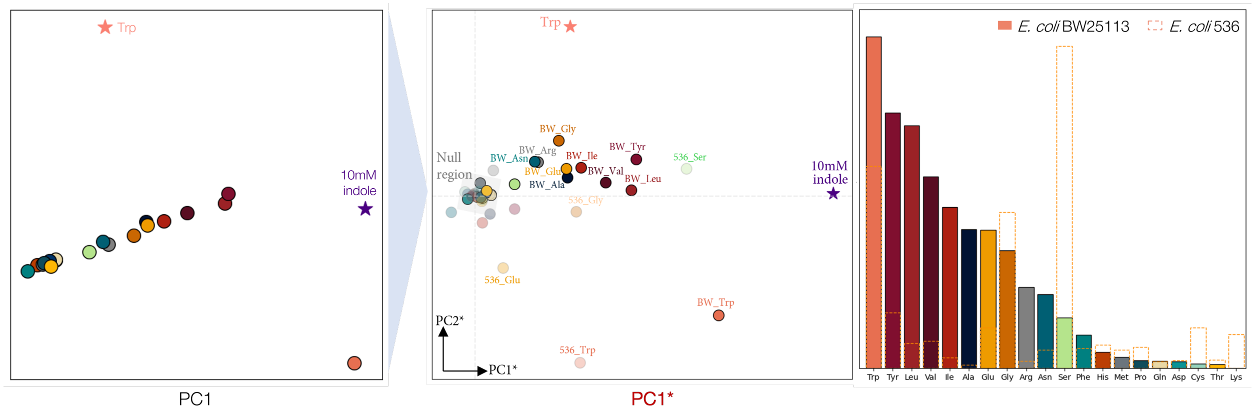


**Supplementary Fig. 3.6: Rotated PCA of supernatant WT minus KO spectra for both *E. coli* BW25113 and 536 strains, as shown in Fig. 3d)**. Left-most plot shows an example of the reference PCA used for the rotation. A null region is defined and both axes rotated until the pure indole lies along the x-axis. Distance in this space from pure indole is therefore measured by PC1*, the new rotated axis.

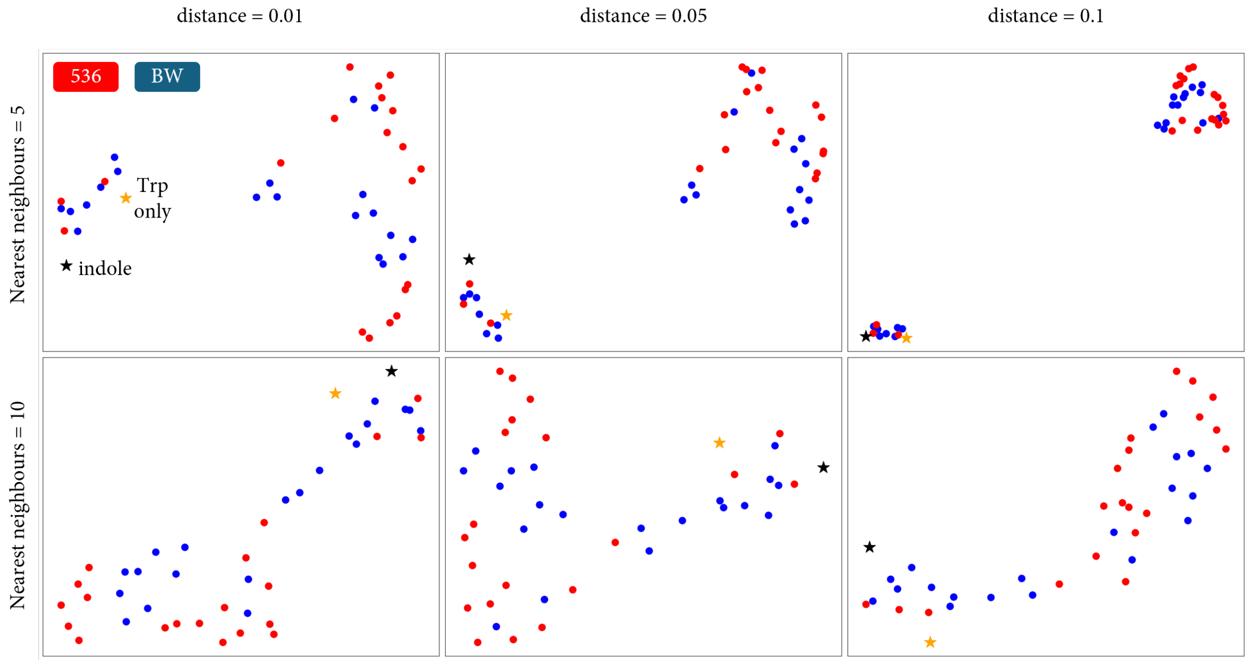


**Supplementary Fig. 3.7: UMAP of WT minus KO spectra for *E. coli* BW25113 and *E. coli* 536 assays, coloured by strain.** Alternative dimensionality reduction techniques such as UMAP show clustering but not a clear projection as with PCA, with further parameter optimisation (distance, nearest neighbours) required before clustering can be obtained. This makes UMAP a less ideal dimensionality reduction technique than PCA for spectral data in this context.

**LC-MS results identifying TnaA-dependent metabolites in *E. coli***

Statistical analysis of the mass spectrometry data shows that ∼60 metabolites are needed to explain 50% of the variance and ∼200 metabolites to explain 90% of the variance (SI Fig. 4.1 (a)), implying a lack of any coherent metabolic pattern across amino acids in marked contrast to the picture presented by the SERS results. This is further highlighted by the distribution of the LC-MS data after PCA, which shows no clustering or pattern.

We analyse the $\Delta$MS peak area changes to locate TnaA-dependent metabolic activity across all 914 metabolites detected (SI Fig. 4.1). Considerable variability is observed, both across strains for the same amino acid (e.g. Trp), and across amino acids for the same strain (e.g. Arg and Ser). Generally, *E. coli* 536 exhibits more TnaA-dependent activity compared to *E. coli* BW25113, with higher $\Delta$MS particularly in the unannotated range. This can be seen more clearly in SI Fig. 4.1b, which shows the number of metabolites above different $\Delta$MS thresholds. Surprisingly, more TnaA-dependent metabolic activity is observed for Cys in $\Delta$MS while both WT and KO show minimal activity in the supernatant and enzymatic SERS assays.

We also determine the degree to which the I* spectral signature, detected in Fig. 2, is a combination of many unknown metabolites. SI Fig. 4.1 (c) plots the rotated PC1* (indole-like) component against the number of unknown TnaA-dependent metabolites identified in $\Delta$MS. Generally, Trp, Tyr and Leu assays (with largest PC1* values) have fewer unknown metabolites compared to assays with smaller PC1* components. Lower PC1* scores require more metabolites, suggesting the indole-like signature is comprised of one or fewer metabolites.

Some of the common metabolites between the two strains (SI Fig. 4.1d, top panel) and between amino acids (bottom panel) are illustrated for metabolites with $\Delta\text{MS}>1.5$ (when WT is 3x higher than KO). At this threshold, for Trp, indole, $\gamma$-aminobutyric acid other six other unknown metabolites are shared between strains. *E. coli* BW25113 when supplemented with the amino acid Phe, yields 52 unique metabolites not present in cultures supplemented with Tyr or Trp, but shares ten metabolites with Tyr and not Trp, and five metabolites with Trp and not Tyr. Metabolites shared across all three aromatic amino acids include trimethylamine N-oxide, a clinically important metabolite found to play a role in cholesterol metabolism, and glycylproline, a metabolite released during the breakdown of collagen.

a)

| Log2 fold-change | 1 | 2 | 3 | 4 | 5 |
|---|---|---|---|---|---|
| *WT vs KO* | *2x higher* | *4x higher* | *6x higher* | *8x higher* | *10x higher* |
| Trp_BW | 57 | 31 | 19 | 14 | 13 |
| Trp_536 | 149 | 81 | 49 | 25 | 15 |
| Tyr_BW | 56 | 25 | 18 | 12 | 10 |
| Tyr_536 | 19 | 5 | 3 | 2 | 1 |
| Leu_BW | 37 | 15 | 6 | 1 | 0 |
| Leu_536 | 11 | 3 | 2 | 1 | 0 |
| Gly_536 | 24 | 1 | 0 | 0 | 0 |
| Arg_BW | 52 | 27 | 17 | 14 | 7 |
| Ser_BW | 86 | 46 | 29 | 24 | 12 |
| Phe_BW | 117 | 67 | 52 | 35 | 16 |
| Phe_536 | 62 | 15 | 7 | 3 | 0 |
| Cys_BW | 145 | 115 | 86 | 60 | 36 |

b)

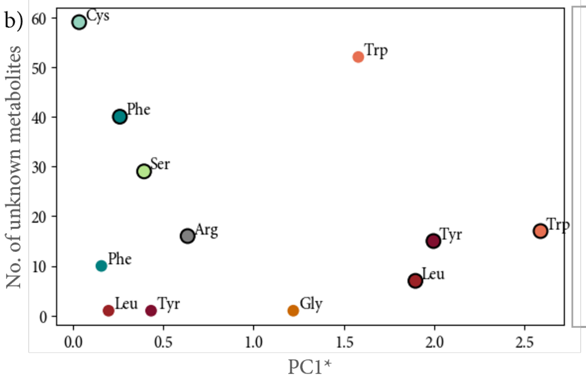


c)

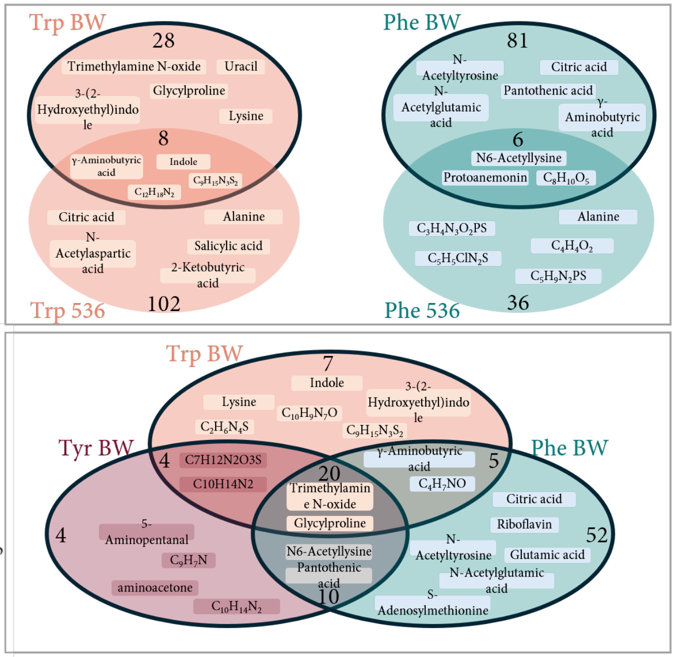


**Supplementary Fig. 4.1: Analysis of LC-MS detected metabolites by log$_2$ fold-change.** (a) Number of metabolites at different fold-changes across selected samples. (b) Plot of PC1* scores (as determined in Fig. 3) against the number of unknown metabolites in each sample, showing generally higher metabolites for assays with a lower PC1* score, indicating the indole-like signature may be comprised of fewer rather than many metabolites. (c) Number of metabolites with a log$_2$ fold-change greater than 1.5 (i.e. 3x higher abundance in WT relative to KO sample) for selected amino acids. Top panel shows intersection across different strains for the same amino acid (Trp and Phe), whilst bottom panel shows intersection across amino acids for the same strain.

**Structural Equation Modelling Analysis of I***

We utilise structural equation modelling (SEM), a multivariate statistical framework to better elucidate the molecular structure of I*. SEM represents linear relationships between observed and latent (unobserved) variables, generalising path analysis wright1934path and confirmatory factor analysis spearman1904general as formalised by joreskog1969general, joreskog1973general. Unlike ordinary least-squares (OLS) regression, SEM can introduce latent constructs inferred from groups of correlated indicators and explicitly models the covariance structure among predictors, allowing correlated residuals to be absorbed rather than collapsed into the error term. A typical SEM analysis defines two layers: a measurement model relating latent variables ($\eta$) to their observed indicators ($X$) via loadings ($\lambda$),

$$X_i = \lambda_i \cdot \eta + \varepsilon_i, \tag{1}$$

and a structural model in which latent and observed exogenous variables jointly predict an endogenous outcome $Y$,

$$Y = \sum_j \beta_j \eta_j + \sum_k \gamma_k X_k + \zeta, \tag{2}$$

with $\beta$ and $\gamma$ the structural path coefficients and $\zeta$ the disturbance. Parameters are estimated jointly by minimising the discrepancy between the observed and model-implied covariance matrices, typically by maximum likelihood under multivariate normality.

**Our use case**

We adapted SEM to answer, given the SERS difference spectrum of I* (bands present, relative intensities, and shifts $\Delta\nu$ relative to free indole), which combination of molecular structural components is most consistent with the observed data? Rather than computing similarity to I* from raw spectral distance or from principal components, the problem was stated as a regression of peak height on molecular structural descriptors and vibrational character of each band, with latent variables representing higher-level structural constructs that are not directly observable. An overview of the analysis workflow is shown in Figure **??**.

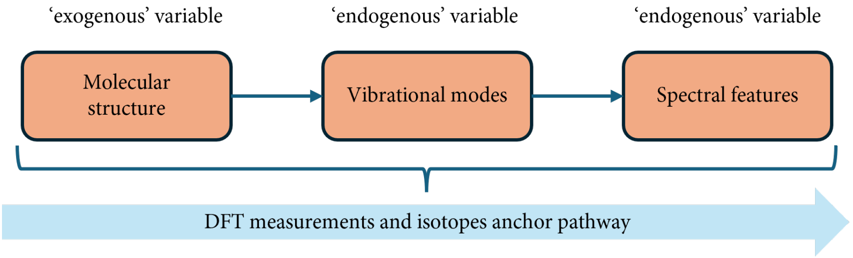


Schematic of our SEM analysis pathway: molecular structure propagates through vibrational mode character to give the observed spectral features. The DFT calculations and isotope-labelled controls anchor each step of this pathway, providing training data from which the path coefficients are estimated.

The SEM was trained on three sources of spectral data. Density functional theory-based spectra (DFT, $n = 7$) for the theoretical anchors of band positions, experimental SERS reference spectra ($n = 9$) of indole and its derivatives (see Fig. 4 in main text), for the experimental anchors, and WT minus KO SERS difference spectra for inverse inference to test the fitted model.

We also created two latent variables: $\eta_{\text{ring},m}$, representing the bicyclic ring system in compound $m$; and $\eta_{\text{oxidation},m}$, representing the extent of ring oxidation. The remaining compound-level descriptors (*e.g.* $X_{\text{C3\_sub}}$, $X_{\text{C2\_oxo}}$, $X_{\text{C3\_oxo}}$, $X_{\text{hydroxyl}}$, $X_{\text{N1\_protonated}}$, $X_{\text{isotope\_d}}$) and all band-level vibrational character descriptors (*e.g.* $X_{\text{ring\_pyrrole}}$, $X_{\text{ring\_benzene}}$, $X_{\text{C-N\_stretch}}$) were retained as directly observed exogenous variables.

For each (compound, band) pair indexed by $(m, k)$, the endogenous variable $Y_{mk}$ is the peak intensity at band $k$ in compound $m$. The structural equation consists of the latent variables, directly observed compound-level descriptors, and band-level vibrational character:

$$
\begin{aligned}
Y_{mk} \; = \; & \underbrace{\beta_1 \eta_{\text{ring},m} + \beta_2 \eta_{\text{oxidation},m}}_{\text{compound via latents}} \\
& + \underbrace{\beta_3 X_{\text{C3\_sub},m} + \beta_4 X_{\text{C2\_oxo},m} + \beta_5 X_{\text{C3\_oxo},m} + \beta_6 X_{\text{hydroxyl},m} + \cdots}_{\text{compound: directly observed}} \\
& + \underbrace{\gamma_1 X_{\text{ring\_pyrrole},k} + \gamma_2 X_{\text{ring\_benzene},k} + \cdots + \gamma_6 X_{\text{C-N\_stretch},k}}_{\text{band-level character}} + \varepsilon_{mk}.
\end{aligned} \tag{3}
$$

Given the fitted coefficients $\hat{\beta}$, $\hat{\gamma}$ and the estimated latent loadings, I*'s observed band heights $\{Y_k^*\}$ were used to solve for the compound-level descriptor vector $\hat{\mathbf{x}}^*$ that best reproduces I*'s spectrum:

$$
\hat{\mathbf{x}}^* \; = \; \arg\min_{\mathbf{x} \geq \mathbf{0}} \|\mathbf{r} - \mathbf{A}\mathbf{x}\|_2^2 \, , \tag{4}
$$

where $r_k = Y_k^* - \hat{\gamma} \cdot \mathbf{x}_k^{(b)}$ is the band-character-corrected residual and $A_{kj} = \hat{\beta}_j$ is the matrix of compound-level path coefficients replicated across all matched bands.

**Supplementary methods**

**Protein purification and active holoenzyme preparation**

The *tnaA* sequence from *E. coli* BW25113 was cloned into pET-19m, to allow for expression of tobacco etch virus protease-cleavable, N-terminally His6-tagged TnaA protein. This pET-19m-tnaA plasmid was electroporated into *E. coli* BL21(DE3) Rosetta, then this strain was inoculated in LB containing 25 $\mu$g/mL of chloramphenicol and 50 $\mu$g/mL of carbenicillin, and incubated overnight at 37°C with shaking. 10 mL of this culture was added to 1 L of LB containing 25 $\mu$g/mL of chloramphenicol and 50 $\mu$g/mL of carbenicillin, incubated at 37 °C with shaking until $OD_{600nm}$= 0.5, then IPTG was added to a final concentration of 1 mM and the culture was incubated overnight at 25 °C with shaking. The following day, the culture was centrifuged at 6000 ×g for 15 minutes at 4 °C, resuspended in 10 mL of ice-cold lysis buffer (50 mM HEPES (pH 7.5), 300 mM NaCl, 5% glycerol), and then a cOmplete™ EDTA-free Protease Inhibitor Cocktail tablet was added. The resuspended cells were then lysed by sonication on ice and centrifuged at 14000 ×g for 30 minutes at 4 °C. The clarified supernatant was then filtered through 0.45 $\mu$m syringe filters and kept on ice.

The filtered supernatant was promptly loaded into a lysis buffer pre-equilibrated 5 mL HisTrap HP column using an AKTA FPLC system. The column was washed with 50 column volumes (250 mL) of lysis buffer, then eluted with 50 mL total volume of a gradient of lysis buffer with increasing proportion of elution buffer (50 mM HEPES (pH 7.5), 300 mM NaCl, 5% glycerol, 300 mM imidazole) and fractions were collected. Fractions with high A280 on the AKTA FPLC were analysed by SDS-PAGE to verify purity and the expected molecular weight of tryptophanase monomers (around 55 kDa). The appropriate fractions were pooled, then loaded into a dialysis bag (30 kDa MWCO) and dialysed overnight at 4 °C against 20 mM HEPES (pH 7.5), 50 mM NaCl, 0.5 mM EDTA, 10% glycerol, 1 mM DTT. The concentration of protein was quantified using Bio-Rad Protein Assay with known standards of bovine serum albumin. The His6 tag was not cleaved, as it was not necessary for subsequent assays.

To ensure that the purified tryptophanase was highly enzymatically active (requiring its cofactor, pyridoxal 5'-phosphate (PLP), and reduced thiol groups [10.1002/9780470122877.ch6]), the dialysed TnaA was incubated with 20 mM DTT for 3 hours at 37 °C, followed by addition of PLP to 25 $\mu$M per 25 $\mu$M of TnaA tetramers (5.5 mg/mL) for an additional 45 minutes at 37 °C, in the dark. To remove DTT that may affect the downstream SERS measurements, the highly active enzyme stocks were passed through PD-10 columns pre-equilibrated with a simple dialysis buffer (0.1 M HEPES (pH 7.8), 10% glycerol), then eluted with the same dialysis buffer. The eluted protein stock was aliquoted, promptly flash frozen in liquid nitrogen, and stored at -70 °C until use.

**Artificial urine media preparation**

Artificial urine media was modified to contain citric acid, sugars, and amino acids at concentrations similar to human urine [1–3]. The recipes for the base and supplement solutions are in Table 1 and Table 2 below.

**Table 1: Composition of the base of artificial urine media, prior to adding supplement solutions**

| Name | Final concentration (mM) |
|---|---|
| Sodium sulfate | 12 |
| Citric acid | 3.3 |
| Trisodium citrate dihydrate | 2.5 |
| Creatinine | 7.8 |
| Urea | 249.8 |
| Potassium chloride | 31 |
| Sodium chloride | 30 |
| Calcium chloride | 1.7 |
| Ammonium chloride | 23.7 |
| Potassium oxalate monohydrate | 0.2 |
| Magnesium sulfate heptahydrate | 4.4 |
| Sodium phosphate monobasic anhydrous | 18.7 |
| Sodium phosphate dibasic anhydrous | 4.7 |
| Uric acid | 1.5 |
| Hippuric acid | 3.5 |
| Thiamine-HCl | 0.003 |

**Table 2: Composition of the supplement solutions of artificial urine media, added to the base recipe in Table 1**

| Name | Final concentration ($\mu$M) |
|---|---|
| L-Tyrosine | 150 |
| L-Galactose | 100 |
| L-Glucose | 500 |
| L-Sorbitol | 100 |
| L-Carnitine | 65 |
| L-Citrulline | 10 |
| L-Ornithine | 65 |
| D-Alanine | 35 |
| D-Asparagine | 15 |
| D-Serine | 125 |
| L-Alanine | 290 |
| L-Arginine | 110 |
| L-Asparagine | 130 |
| L-Aspartic acid | 150 |
| L-Cysteine | 1000 |
| L-Glutamic acid | 100 |
| L-Glutamine | 520 |
| L-Glycine | 1400 |
| L-Histidine | 600 |
| L-Isoleucine | 20 |
| L-Leucine | 40 |
| L-Lysine | 240 |
| L-Methionine | 15 |
| L-Phenylalanine | 90 |
| L-Proline | 15 |
| L-Serine | 310 |
| L-Threonine | 190 |
| L-Tryptophan | 80 |
| L-Valine | 60 |
